# The functional landscape of the human phosphoproteome

**DOI:** 10.1101/541656

**Authors:** David Ochoa, Andrew F. Jarnuczak, Maja Gehre, Margaret Soucheray, Askar A. Kleefeldt, Cristina Viéitez, Anthony Hill, Luz Garcia-Alonso, Danielle L. Swaney, Juan Antonio Vizcaíno, Kyung-Min Noh, Pedro Beltrao

## Abstract

Protein phosphorylation is a key post-translational modification regulating protein function in almost all cellular processes. While tens of thousands of phosphorylation sites have been identified in human cells to date, the extent and functional importance of the phosphoproteome remains largely unknown. Here, we have analyzed 6,801 publicly available phospho-enriched mass spectrometry proteomics experiments, creating a state-of-the-art phosphoproteome containing 119,809 human phosphosites. To prioritize functional sites, 59 features indicative of proteomic, structural, regulatory or evolutionary relevance were integrated into a single functional score using machine learning. We demonstrate how this prioritization identifies regulatory phosphosites across different molecular mechanisms and pinpoint genetic susceptibilities at a genomic scale. Several novel regulatory phosphosites were experimentally validated including a role in neuronal differentiation for phosphosites present in the SWI/SNF SMARCC2 complex member. The scored reference phosphoproteome and its annotations identify the most relevant phosphorylations for a given process or disease addressing a major bottleneck in cell signaling studies.

## Introduction

Protein phosphorylation is a post-translational modification (PTM) that can alter protein function at seconds to minutes time scale. The addition of a phosphate group, mainly to serine, threonine and/or tyrosine, can regulate a protein by changing its conformation, interactors, localization, and degradation rate, among other mechanisms. In fact, this PTM is involved in the regulation of most biological processes and its misregulation has been linked to several human diseases (Lahiry et al. 2010; Torkamani et al. 2008). The full extent of protein phosphorylation events occurring in human cells is still an open question that is under active investigation by using mainly mass spectrometry (MS) proteomics approaches. Over the past few years, this technology has been developed to the point of achieving high accuracy and coverage of phosphosite identification (Aebersold and Mann 2016). Notably, an in-depth study of a single cell type identified over 50,000 phosphopeptides and suggested that 75% of the proteome may be phosphorylated (Sharma et al. 2014). Such studies have led to the identification of a large number of human phosphosites with over 200,000 currently aggregated in the PhosphoSitePlus (PSP) resource (Hornbeck et al. 2019).

Although analytical challenges still remain, the bottleneck in the study of protein phosphorylation is largely shifting from analytical challenges towards its functional characterization (Needham et al. 2019). Given that protein phosphorylation can be poorly conserved and that kinases often have some degree of promiscuous activity, it has been suggested that not all phosphorylation is relevant for fitness (Landry, Levy, and Michnick 2009; Kanshin et al. 2015; Beltrao et al. 2013). Even if all phosphosites have a functional role, it is likely that they are not equally essential for the organism. Therefore, prioritization strategies are crucial to facilitate the discovery of highly relevant phosphosites (Beltrao et al. 2012). Different methodologies have been proposed for such prioritization, that including e.g. identifying phosphosites that are highly conserved (Studer et al. 2016), those located at interface positions (Šoštarić et al. 2018; Betts et al. 2017; Nishi, Hashimoto, and Panchenko 2011), showing strong condition-specific regulation, or near other PTM sites (Beltrao et al. 2012). Phosphosite mutational studies followed by experimental phenotypic assays have been used to identify functionally relevant phosphorylations in limited cases, but cannot yet be easily applied to the study of human phosphorylation at scale.

Machine learning based approaches have been used to rank functional elements according to functional relevance including, for example, gene essentiality (Seringhaus et al. 2006). This remains an unexplored approach to study the functional relevance of protein phosphorylation in a comprehensive fashion. Here, we generated a state-of-the-art human phosphoproteome by jointly analyzing 6,801 publicly available phospho-enriched MS proteomics experiments, identifying 119,809 human phosphosites. For each detected phosphosite, we compiled annotations covering 59 features that could be indicative of biological function and integrated them into a single score of functional relevance, named here the phosphosite functional score. This score can correctly identify regulatory phosphosites for a diverse set of mechanisms and predict the impact of mutations in phosphosites. We provide experimental evidence for novel regulatory sites including a role in neuronal differentiation for phosphosites present in the SWI/SNF SMARCC2 complex member.

## Results

### Mass spectrometry-based proteomics map of the human phosphoproteome

In order to develop an integrated approach to characterize phosphosite functional relevance, we first set out to create a state-of-the-art MS-based definition of the human phosphoproteome. In order to obtain a comprehensive phosphoproteome, we manually curated 112 human public phospho-enriched datasets derived from 104 different cell types and/or tissues from the PRIDE database **(Table S1)** (Vizcaíno et al. 2016). Next, we jointly re-analyzed using MaxQuant the subset of 6,801 human MS experiments passing the quality control criteria, corresponding to 575 days of accumulated instrument time (**Methods**) (Cox and Mann 2008). The joint analysis (deposited in PRIDE, dataset PXD012174) ensured an adequate control of the false discovery rate, preventing the accumulation of false positive identifications in parallel searches (**Methods**). As a result, we identified a total of 11.7 million phosphorylated peptide-spectrum matches (PSM-level FDR < 1%), corresponding to 181,774 phosphopeptides spanning a total of 203,930 phosphorylated serines, threonines or tyrosines. Of these, only 119,809 phosphorylated sites passed the 1% site-level FDR correction (59% true positive sites). 90,443 of those sites were classified as Class I (Olsen et al. 2006). The low percentage of true positive sites suggests that the accumulation of phosphosite identifications from multiple independent searches - as archived in phosphosite databases - might be strongly enriched for potential false positives if not adequately corrected at an aggregated level. The heterogeneity of biological samples analyzed in this study facilitated the identification of phosphosites present in proteins expressed in a wide range of tissues and cell types, healthy individuals and tumor samples (**Figure 1a**). For example, we could identify a large number of tissue-specific phosphosites, including phosphorylation of myosin in muscle cells, gastric mucin in stomach samples or melanin in healthy skin samples (**Figure 1b**).

**Figure 1.**
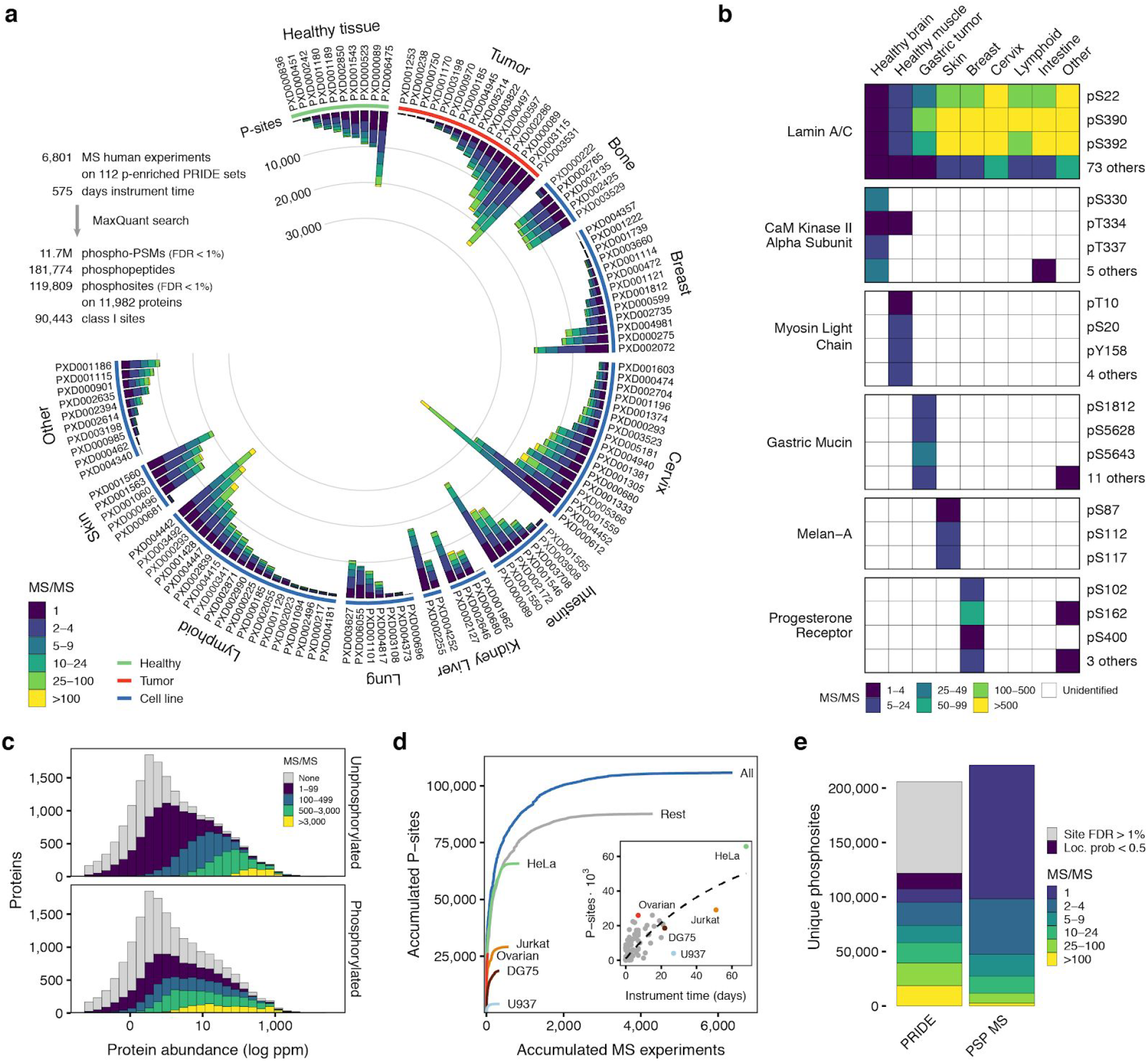
Comprehensive catalog of in-vivo human phosphosites. a) Number of phosphosites (Localisation probability > 0.5) binned by the number of peptide-spectrum matches coming from the re-analyzed human phospho-enriched datasets curated from PRIDE. b) Examples of broad or tissue-specific phosphosites with spectral count information. c) Phosphopeptide and unphosphorylated peptide MS/MS support for all human proteins binned using the consensus protein abundance from PaxDb d) Cumulative increase in the number of phosphosites for each added MS experiment by biological origin: all samples (blue), top 5 most common cell lines or tissues or remaining samples (grey). MS experiments were sorted by size. Inset - Accumulated instrument time and the total number of phosphosites identified per sample. e) Total number of unique identified phosphosites and MS/MS support in the combined PRIDE analysis and PSP.

To evaluate the phosphorylated proteome coverage, we studied the proportion of phosphoproteins when stratifying all proteins according to their consensus overall abundance (Wang et al. 2015). From the 14,154 reviewed UniProt proteins identified, 11,982 (85%) contained at least one FDR-corrected phosphosite. While we observed a bias towards the identification of more abundant proteins, the trend is similar to that of the non-modified peptides present in the samples (**Figure 1c**). This ratio of phosphoproteins remained constant regardless of the reported protein abundance in PaxDb (**Figure S1**) and is similar to previous findings (Sharma et al. 2014). While there are some cell types that are very commonly represented in the re-analyzed datasets, such as HeLa cells, the identified phosphosites are not strongly biased by these common samples. The exclusion of phosphorylation information retrieved from the five most studied cell types (31% of the total instrument time), still results in 83% of the total identified phosphosites (**Figure 1d**). Together, these results suggest that we have achieved very high coverage of phosphosite identification.

For benchmarking purposes, we compared the identified phosphoproteome against the 221,236 human phosphosites reported by MS in the PSP database (January 2018) (**Figure 1e**) (Hornbeck et al. 2019). While 11.5% of the high-confidence sites in our study are supported by only 1 MS/MS evidence (mass spectrum), 55% of the PSP sites have this level of support. In absolute numbers, we identified 73,973 phosphosites supported by 5 or more MS/MS evidences, while only 47,448 sites are supported by the same number of evidences in PSP. These results point to the importance of aggregating the growing body of MS phosphoproteomic data while maintaining the statistical reliability.

### Prioritizing functional human phosphosites

Having identified a comprehensive high confidence human phosphoproteome, we then calculated for every phosphosite a diverse set of features that could indicate importance for fitness (**Methods**). These properties can be grouped based on whether they add functional support due to the MS-evidence, the phosphosite regulation (Ochoa et al. 2016), the structural environment, or the evolutionary conservation. For each group, multiple features attempt to detect different functional characteristics. For example, evolutionary conservation was captured by independent features quantifying residue conservation (Vaser et al. 2016), spatial conservation of the phosphorylation within protein domain families (Strumillo et al. 2018) and the phosphosite evolutionary age, expanding to multicellular species a phylogenetic-based approach previously described for fungal species (Studer et al. 2016). Altogether, we calculated a set of 59 features that were used to annotate all phosphosites (**Methods and Table S2**). We illustrate the value of some of these features for identifying regulatory sites in the 65-80 amino acid region of MAF1, a protein involved in the mTORC1 signaling pathway (**Figure 2a**). While the phosphorylations pS68 and pS75 are known to inhibit the MAF1 RNA pol III repression function, three other MS-identified phosphosites in the region have currently unknown functional roles (Michels et al. 2010). Relative to the other sites, the two known regulatory sites showed a higher number of spectral counts and are observed across a larger number of samples. Additionally, the phosphorylations are more conserved across species, more likely to match the substrate specificity of kinases and show condition-specific regulation, including downregulation when treated with the mTOR inhibitors rapamycin or Torin1.

**Figure 2.**
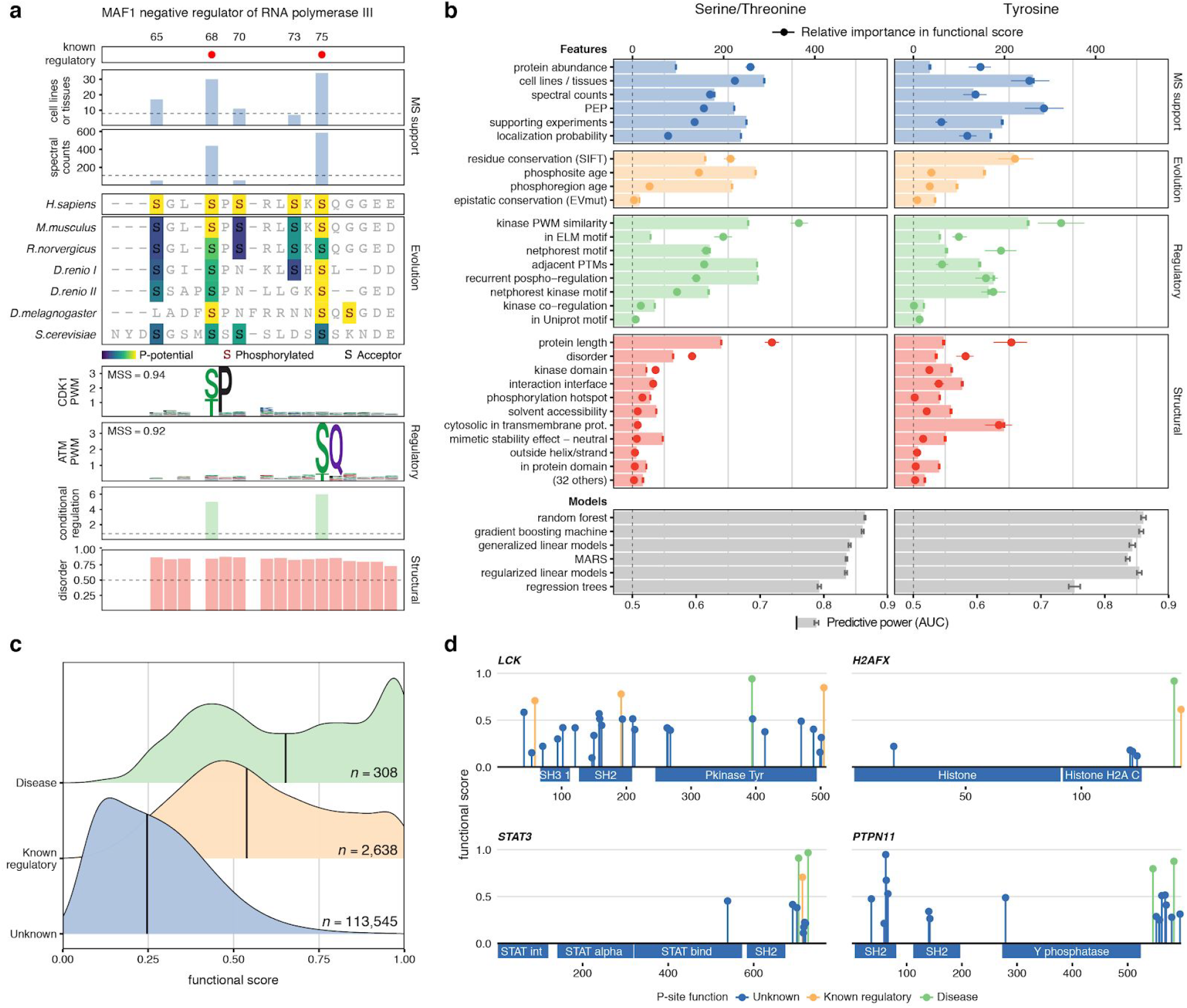
A functional score for human phosphosites. a) MAF1 phosphosites in the 65-80 region with feature annotations b) Feature discriminative power (AUCs) after repeated cross-validation between phosphosites of known and unknown function (AUC, red, green, yellow and blue bars). Discriminative power (AUCs) after integrating all features using different machine learning algorithms (AUC, grey bars). The contribution of each feature to the final model (point-ranges). c) Distribution of functional scores for all phosphosites (blue), with known regulatory roles (yellow) and associated with human diseases (green). d) Functional score and regulatory function for phosphosites in different protein families.

Benefiting from the orthogonal nature of some of these attributes, we next sought to integrate a set of 59 normalized features into a single score that would prioritize those phosphosites most relevant for fitness. We used 2,638 phosphosites curated by PSP and known to regulate protein function to discriminate, using machine learning, the distinctive properties of functional serine and threonine phosphorylation (S/T) and tyrosine phosphorylation (Y), separately. We first asked how well each of the 59 features independently predicted the known regulatory sites. In other words, how a feature (e.g. conservation) could differentiate the known regulatory sites - true positives - from the background of all sites - bars in **Figure 2b**. The most informative features for function included the number of different cell lines or tissues in which the site had been identified, the phosphosite age, how well it matched a kinase specificity model, how often the phosphosite was regulated in a panel of different perturbations and the presence of neighboring PTMs.

Different machine-learning algorithms were then tested for their capacity to integrate the 59 features, in order to find the best combination of signals that could characterize the known regulatory sites (**Methods**). The top 5 algorithms had overall similar performance after hyperparameter optimization and repeated cross-validation as judged by the area under the Receiver Operating Characteristic curve (AUC). The final model selected was a gradient boosting machine that achieved an average AUC of 86.1% and 85.7% for ST and Y, respectively (Friedman 2002). The contribution of the most relevant features once included in the integrative model is shown as point ranges in **Figure 2b** and denote the relevance of each feature when combined with the rest of the features. For example, the consensus protein abundance significantly increased its relevance when integrated with other features, indicating protein abundances are not good predictors of function by themselves, but facilitate the interpretation of other features within the model.

The final model - named here phosphosite functional score - was used to generate a score for each of the MS-identified phosphosites, reflecting their importance for organismal fitness (**Table S3**). In **Figure 2c,** we show how this score ranked known functional phosphosites higher than the overall background of all other phosphosites and, more interestingly, phosphosites important for human disease - information not included in the model - ranked higher than the other two sets. The functional score could also correctly distinguish relevant phosphosites across different protein families as illustrated in four different examples, including the LCK kinase, the STAT3 transcription factor, the PTPN11 phosphatase and the H2AFX histone (**Figure 2d**). In all cases, the phosphosites of known function or associated with diseases were among the top-ranked. Extensive literature search also pointed to highly scored phosphosites that, despite not being included in the true positive set derived from PSP, had supporting evidence of known regulatory function (**Figure S2**). For example, valosin-containing protein (VCP) pY805 is ranked as the highest scoring (0.65) phosphosite in the protein and it’s known to disrupt interaction with both PNGase and Ufd3 (Zhao et al. 2007). Similarly, alanine or glutamate mutation of the best-scored S/T site pS6 (0.81) in SDCBP abolished interaction with ubiquitin, as demonstrated by His-Ub pulldown assays (Rajesh et al. 2011). Phospho-mimicking mutations in the highly scored p62/SQSTM1 pS24 (0.72) restored polymerization instead (Christian et al. 2014). The phosphosite functional score recapitulated all this knowledge in spite of their absence from the true positive list of functional phosphosites used to derive the model.

### Identifying functional phosphosites involved in diverse mechanisms

Phosphorylation modulates protein function via several different molecular mechanisms. The distributions of phosphosite functional scores for annotated regulatory sites suggest the method is not strongly biased towards specific mechanisms, detecting with similar accuracy regulatory phosphosites relevant for protein degradation, localization, conformation, interactions, enzymatic activity, and other functions (**Figure S3**). Next, we explored the potential of the phosphosite functional score to prioritize known and novel regulatory functions related to protein-protein interactions and transcriptional activity.

The presence of the phosphosite in the structural model of a protein interaction interface is one of the features used for the generation of the functional score. In our dataset, we identified phosphosites with a high functional score that were located at protein interface models, as being likely to regulate protein-protein interactions (**Methods,** provided in **Table S2**). For example, a low-abundance phosphosite Y34 of the Ras Homolog Family Member A (RHOA) had an unknown function according to PSP and ranked as a highly functional site (0.56) partly due to its presence in multiple interaction interfaces. Indeed, it has been reported that mutation of Y34 to asparagine (Houssa et al. 1999) or phenylalanine (Worby et al. 2009; Uezu et al. 2012) can disrupt certain interactions. Altogether, these findings suggest the phosphorylation at this position could shift the interaction affinity (**Figure S4**). Independently, we sought to validate as a proof-of-principle a candidate regulatory phosphosite in a protein-protein interface with no previous experimental evidence. The PLK1-regulated S60 is the best scoring phosphosite (0.65, Figure 3a) of the RAN Binding Protein 1 (RANBP1) (Hwang, Ji, and Jang 2011). Some of the most relevant features of pS60 are its strong upregulation (logFC 4.44) under okadaic acid treatment and its proximity to the Ran interaction interface and the transmembrane nuclear transporter Ran GTPase Activating Protein 1 (RanGAP) (**Figure 3b**, PDB:1k5g). In order to evaluate the potential of pS60 to regulate RanBP1 interactions, we performed a pull-down experiment followed by MS on the 3xFLAG tagged RANBP1 WT and quantitatively compared it with RANBP1 S60E (**Methods**, **Table S4**). Among the top scoring interactors, proteins RAN, RCC1 and NEMP1 were found. The phospho-deficient mutant binding remained similar to the WT in the case of RAN and RCC1 but showed a reproducible decrease (p < 0.003) in binding capability to NEMP1, an interaction partner previously described in other eukaryotes (**Figure 3c**) (Shibano et al. 2015).

**Figure 3.**
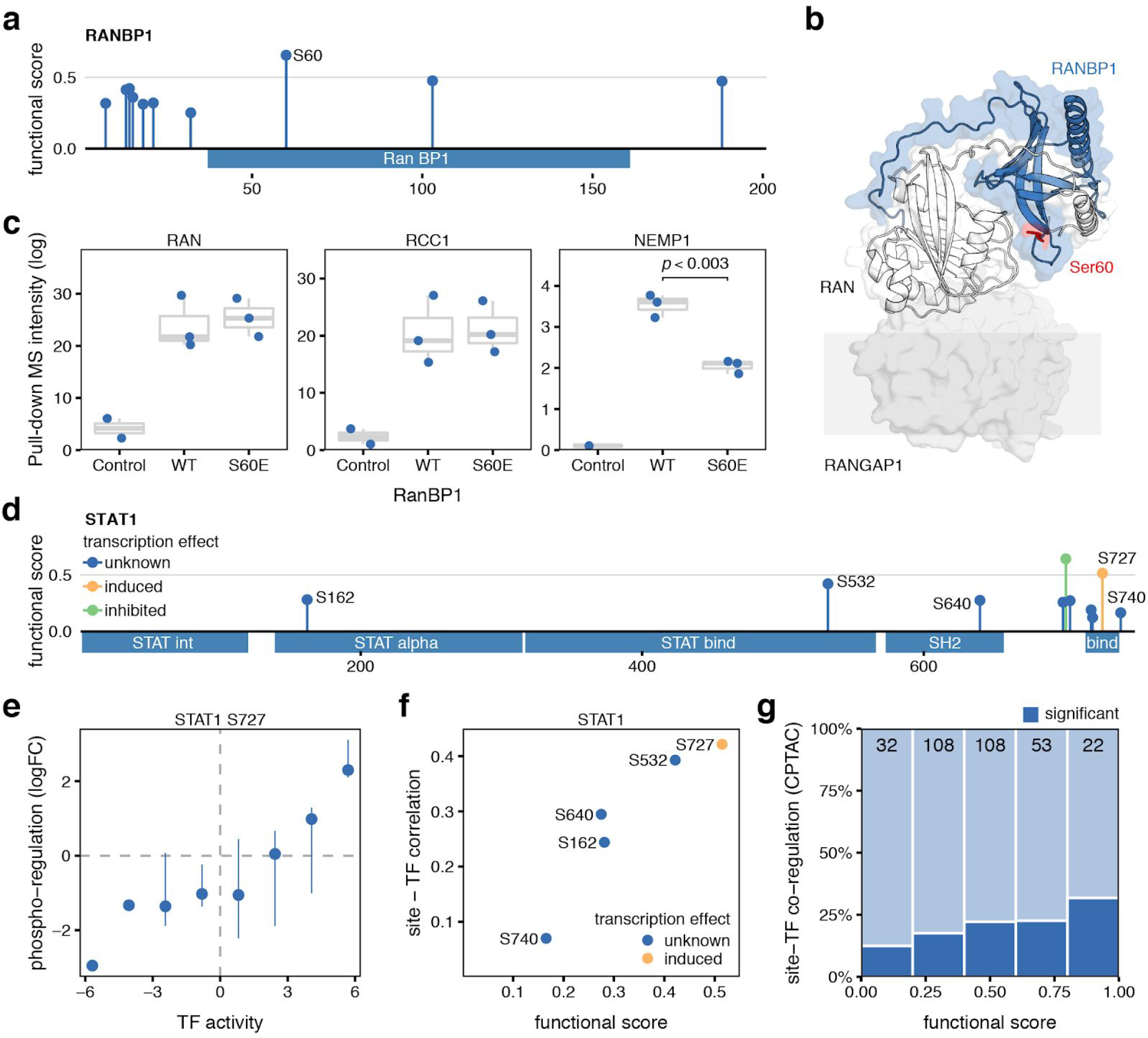
Identification of functional sites regulating protein interaction and transcriptional activity. a) Functional score for phosphosites identified in RANBP1. b) Structural model of RANBP1 in complex with RAN (PDB:1k5g). c) MS binding quantification for RANBP1 interaction partners (RAN, RCC1, and NEMP1) pulling-down the control, the WT, or the S60E RANBP1 mutant. d) Functional score of phosphosites identified in STAT1 with known activating (yellow) and inhibitory (green) regulatory activity. e) Correlation between the changes phosphorylation levels of the known activation site of STAT1 (S727) with the changes in estimated STAT1 transcriptional activity. f) Relationship between STAT1 phosphosite functional score and their correlation with activity. g) Fraction of phosphosites in TFs showing significantly correlated changes with the corresponding TF activity, stratified by their functional score. Data obtained from TCGA and CPTAC consortia (see Methods).

Along with protein-protein interactions, we explored how the phosphosite prioritization could be used to identify sites implicated in the regulation of transcriptional activity. Changes in the activation state of transcription factors (TFs) across different samples can be approximated by quantifying the changes in expression of their known target genes (Garcia-Alonso et al. 2018). By analyzing 77 breast cancer samples where phosphorylation (Mertins et al. 2016) and gene expression (Cancer Genome Atlas Network 2012) had been collected from the same tumor, we correlated the changes in phosphosite levels within TFs with the changes in estimated TF activity (**Methods**). We exemplify this approach with phosphosites in STAT1 (**Figure 3d**), where pS727 is necessary for full transcriptional activation (Wen, Zhong, and Darnell 1995). Across samples, we observed as expected that increased phosphorylation of S727 was associated with higher estimated TF activities (**Figure 3e**). Interestingly, across all STAT1 phosphosites, the phosphosite functional score stratified all quantified sites based on the correlation between the phosphorylation levels and the TF activity (**Figure 3f**). This prioritization of sites is particularly relevant in cases such as STAT1, where the phosphorylated protein has been reported as a positive prognostic marker for breast cancer (Widschwendter et al. 2002; Tymoszuk et al. 2014). We expanded this analysis to the 371 phosphosites in 82 TFs with available paired data and good quality regulons. Although the set of sites with significant site-TF correlation is relatively small (19%), we observed that the functional score was able to prioritize sites in which the changes in phosphorylation strongly correlated with the TF activity (**Figure 3g**).

### Impact of genetic variation on highly functional phosphosites

A functional phosphosite is expected to introduce a genetic constraint in the genome at that position. Highly functional phosphosites should, therefore, present a different tolerance to variation compared to lower functional ones. To demonstrate this, we analyzed for every phosphosite the available information on allele frequency of variants in natural populations (**Figure 4a**) (Lek et al. 2016) and the clinical significance of mutations on human diseases (**Figure 4b**, **Methods**) (Landrum et al. 2018). In both cases, the higher the functional score the higher the chances that a missense mutation causes a severe consequence. We found that mutations in phosphosites with a high functional score were more likely to be rare in human populations (**Figure 4a**) and more likely to be pathogenic (**Figure 4b**). Overlying this information coming from known disease-causing mutations with the annotated phosphoproteome provides an opportunity to further characterize the underlying mechanisms of these genetic variants. For example, the S172P mutation in Tubulin, Beta 2B (TUBB2B) has been associated with polymicrogyria, a cortical developmental disease causing poor incorporation of tubulin into the microtubules (Jaglin et al. 2009). The finding of a high functional phosphosite in S172 (0.43) as one of the best scoring phosphosites in the protein (max 0.45), not only underscores the functional importance of this position (**Figure 4b**) but also implicates phospho-regulation as an important disease-related mechanism. Associating disease variants with the functionally annotated phosphoproteome can facilitate the disease interpretation in a signaling context, expanding the possibility for diagnostic and therapeutic strategies.

**Figure 4.**
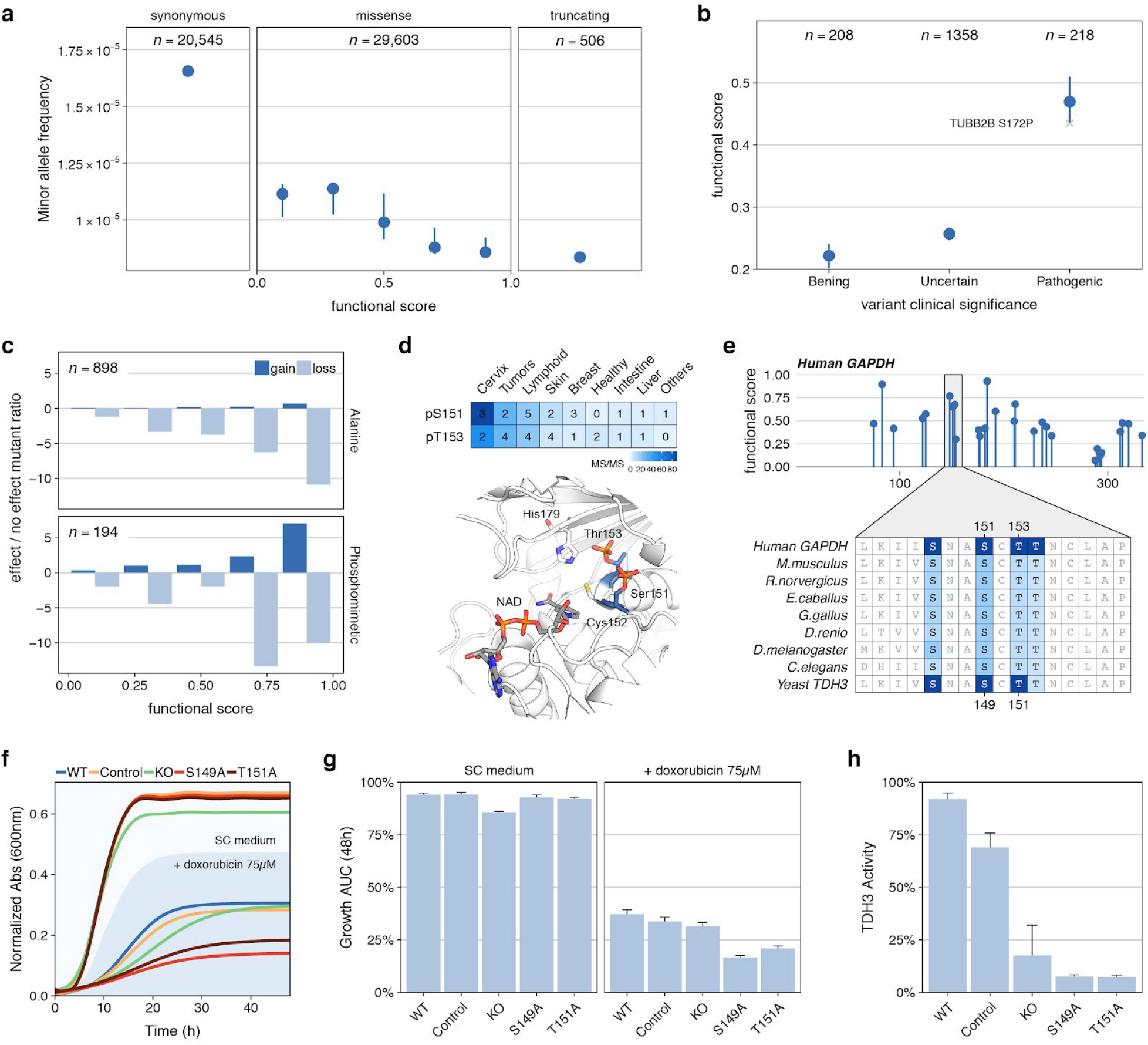
Consequences of genetic variants for phosphosites with high functional scores. a) Median (CI > 95%) minor allele frequency for variants sorted by phosphosite functional score and compared with synonymous and stop codon causing variants occurring at phosphosite positions. b) Mean functional score (CI > 95%) for phosphosites at positions with mutations found in patients and having benign, uncertain or pathogenic consequences. c) Fold ratio of mutations in phosphorylated positions reported in mutagenesis studies having gain/loss of function effects versus no effect and stratified by functional score. d) MS evidence for the phosphorylation of S151 and T153 in GAPDH and their structural context flanking a catalytic cysteine e) Position and functional score for all GAPDH phosphosites and alignment of two human phosphosites (S151 and T153) to the corresponding S. cerevisiae TDH3 (S149 and T151). f) Consensus growth curves and (g) mean and standard error of the area under the growth curve for wild type (WT), control, GAPDH knockout (KO) and S149 and T151 phospho-deficient mutants in the presence or absence of doxorubicin (75 µM). h) TDH3 activity measured in extracts obtained from control and mutant strains.

The introduction of mutations in highly scored acceptor residues as a result of mutagenesis studies is also expected to cause important functional consequences. To systematically study this effect, we curated information on protein gain/loss of function as a consequence of 1,092 mutations in phospho-acceptor residues (**Methods**, **Figure 4c**). In line with our expectation, the higher the phosphosite functional score the higher the chances a mutation causes an effect (**Figure 4c**). Interestingly, mutations of highly scored acceptor residues to alanine were predominantly found to cause a loss-of-function effect, while mutation to negatively charged residues that could mimic the phosphorylation had also an increased chance of causing a gain-of-function effect. To further expand on this idea, we selected to study two highly scored phosphosites of unknown function in glyceraldehyde 3-phosphate dehydrogenase (GAPDH). The two sites selected - S152 and T153 phosphorylation - flank the catalytic cysteine C152 in a highly conserved region (**Figure 4d**). Phosphorylation was consistently identified by MS and their phosphosite functional scores (0.63 and 0.70, respectively) indicated a potential regulatory function (**Figure 4e**). Based on the phosphosite functional scores and the additional features, we hypothesized these phosphosites to be important for the regulation of GAPDH enzymatic activity and functionally conserved between human and yeast.

To validate this hypothesis, we created two phospho-deficient strains - S152A and T153A - in the conserved *S. cerevisiae* main GAPDH gene (TDH3) and measured their growth together with the TDH3 gene knockout (KO) in the presence and absence of the topoisomerase inhibitor doxorubicin **(Methods)**. Deletion of TDH3 has been previously reported to cause slow-growth under doxorubicin treatment in a systematic screening, a phenotype corroborated in our assay (Westmoreland et al. 2009). Interestingly, more acute growth defects were observed for the 2 phospho-deficient strains when treated with doxorubicin **(Figures 4f and 4g)**. TDH activity assays in protein extracts under no treatment pointed to a depleted enzymatic activity of the 2 phospho-deficient strains **(Figure 4h)**. Both the growth rate under doxorubicin and the enzyme activity of the point mutants were lower than the observed for the KO strain suggesting that the TDH3 KO may have some compensation via TDH1/TDH2 that is not seen in the mutants. Alternatively, the mutants may have a dominant negative effect over TDH1/TDH2. Although based in only these observations, the results indicate that the phosphosite functional score of highly conserved phosphosites could potentially be extrapolated to other organisms.

### Novel regulatory phosphosites in the hSWI/SNF remodeling complex member SMARCC2

The human phosphoproteome and the phosphosite functional score serve as a resource for the prioritization of novel regulatory sites that might control particular cellular processes. Here, we exemplified the usefulness of the functional score to interrogate regulatory sites governing the function of SMARCC2/BAF170 protein during neuronal differentiation. As part of the SWI/SNF chromatin remodelling complex, SMARCC2 is known to play an important role in neurogenesis (Tuoc et al. 2013) with studied gene mutations previously associated with autism (Neale et al. 2012). SMARCC2 expression is regulated during the differentiation of embryonic stem cells towards the commitment to neuronal precursors. During this transition, SMARCC2 replaces one of the copies of SMARCC1 in the neural progenitor-specific complex hSWI/SNF complex (also known as npBAF complex). The introduction of SMARCC2/BAF170 into the npBAF complex directly recruits the REST (RE1-silencing transcription factor-corepressor) complex an interaction that is important for neurogenesis (Tuoc et al. 2013). We identified 2 high scoring phosphosites in SMARCC2 within the hyper-phosphorylated SWIRM-associated domain (**Figure 5a**). We hypothesize that S302 and S304 might have an important regulatory function during neurogenesis.

**Figure 5.**
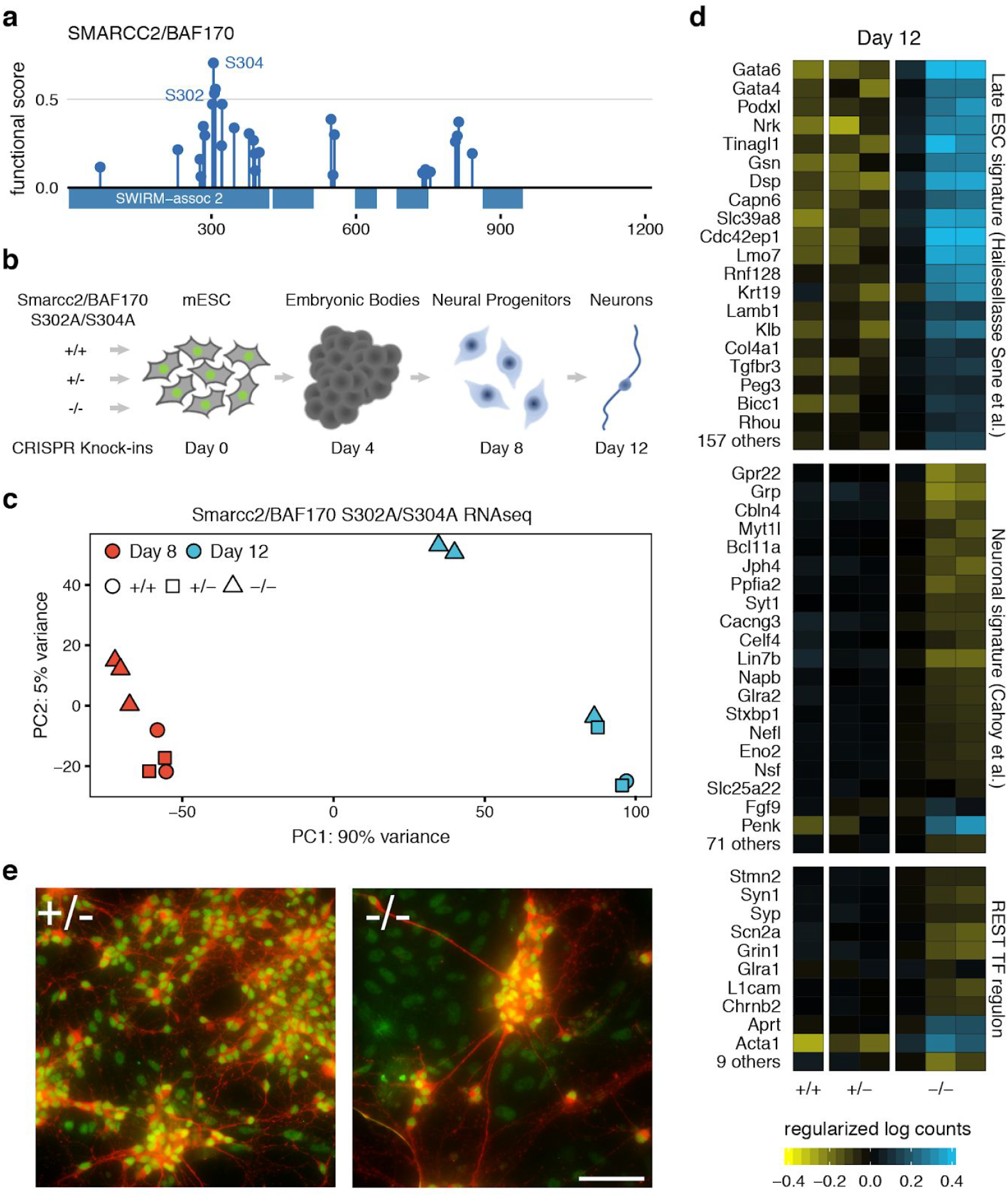
Smarcc2 S302A/S304A homozygous mutants show delayed neuronal differentiation. a) Functional score for SMARCC2/BAF170 phosphosites. b) Design for CRISPR Knock-in mutagenesis of control (+/+), heterozygous (+/-) and homozygous Smarcc2/Baf170 S302A/S304A mutation followed by expected neuronal differentiation timeline. c) PCA plot of gene expression obtained from RNA-seq of the clonal lines. d) Normalized RNA-seq log counts of genes contained in signatures of Late ESC, neurons or transcriptionally regulated by REST. e) Merged immunofluorescence images of differentiated cells on day 12 stained with antibodies against neuronal microtubules Map2 (red) and Smarcc2/Baf170 (Green). Scale bar represents 25 µm.

To study the functional role of the selected SMARCC2 phosphosites, we made use of a murine model of neuronal differentiation and gene knock-in system (**Figure 5b**). After verifying the two phosphosites are identified by MS in neuronal tissues (Liu et al. 2018), we used CRISPR-based mutagenesis in mouse embryonic stem cells (mESC) to independently generate 3 homozygous and 2 heterozygous clones of the double alanine mutant S302A/S304A. For every clone, neuronal differentiation was subjected over a period of 12 days. In agreement with previous findings (Tuoc et al. 2013), an increase of the Smarcc2 protein expression was observed by day 12 (**Figure S5**), concurrent with the formation of mature neurons (**Figure S6**). Based on this timeline, RNA-seq analysis was performed at days 8 and 12 of differentiation, corresponding to the neuronal progenitor cells and mature neurons, respectively. When comparing the mRNA levels using principal component analysis (PCA, **Figure 5c**), the independent clones showed a strong agreement in their transcriptomics profiles at each time point. The major driver of changes (PC1, **Figure 5c**) in the gene expression profile was time, with 8-day samples clearly separated from the 12-day ones. The second major driver (PC2, **Figure 5c**) was the mutational status, with the homozygous clones showing separation from the heterozygous and CRISPR controls. Although the three homozygous clones were separated by PC2 from the rest of genetic backgrounds, one of the homozygous clones did however show a closer gene expression to the heterozygous - CRISPR control group. Overall, these results suggest that the heterozygous mutants have a gene expression state that is not different from the CRISPR control. On the contrary, the homozygous mutations have a clear effect on gene expression, which is higher after 12 days. A linear model identified 4,776 genes (p < 0.05) showing a gene expression difference that was only dependent on the mutational status. This significant transcriptional difference is compatible with a change in the differentiation process as a consequence of the SMARCC2 mutations.

Further analysis of the temporal transcriptomics indicated that the homozygous mutants at day 12 displayed overall transcriptional similarities with the 8-day clones, suggesting some delay in the differentiation process affecting the phospho-deficient clones. Moreover, the homozygous mutants showed upregulation of gene signatures associated with murine pluripotency (Hailesellasse Sene et al. 2007) and significant downregulation in murine signatures associated with neuronal differentiation (Cahoy et al. 2008), as compared to the heterozygous and CRISPR control clones (**Figure 5d**). In addition, a large proportion of the known target genes of RE1-Silencing Transcription factor (REST) - a transcriptional repressor of neuronal genes in non-neuronal tissues - were significantly down-regulated in the homozygous mutant (**Figure 5d**). Neuronal morphology also showed less differentiation in two homozygous clones, which is consistent with the gene expression results. At 12 days, we observed cell aggregates in the homozygous mutant with fewer neurites and numerous Smarcc2 marked cells without the neuronal marker (e.g., Map2) (**Figure 5e**). Overall, these results point to a significant delay in the differentiation in the homozygous clones, suggesting that these phosphorylation sites play a role in the regulation neuronal differentiation by SMARCCs2.

## Discussion

Benefiting from the unprecedented amount of proteomics data generated by the community, we have re-analyzed a substantial number of phospho-enriched MS proteomics datasets deposited in the PRIDE database, in order to generate a comprehensive and state-of-the-art human phosphoproteome. The aggregation of diverse cell lines and tissues, together with heterogeneous perturbations and experimental designs, have facilitated the broad identification of phosphosites including sites in several tissue-specific proteins. The large fraction of phosphosites identified when excluding the most studied cell lines (e.g. HeLa cells) stresses the robustness of this phosphoproteome and the high coverage achieved. However, some limitations need to be taken into consideration. The difficulties to detect certain phosphopeptides - like those in low abundant proteins - or to accurately localize phosphorylation events, point to a still incomplete phosphoproteome. Rarefaction curve analysis (**Figure S7**) of the analyzed samples also suggests we have not yet reached saturation on the number of identified sites and the upper limit of phosphosites remains unknown. Our analysis makes clear that the inadequate aggregation of parallel MS searches can lead to very substantial accumulation of false positives which may be the case for public resources. This can result in a large overestimation of the total number of phosphosites expected for the human phosphoproteome. Our priority has been to provide a high confidence list of phosphosites, implying that some good quality sites might not have made the final list due to the stringent criteria. The scale and level of curation of the spectral data analyzed (PRIDE dataset PXD012174) constitute an unprecedented resource to further develop new methods and strategies necessary to understand phosphoproteomics at large scale.

For these many phosphorylation events identified here, we have devised an approach to prioritize functional phosphosites by integrating indicative features using machine learning. The functional score has recapitulated phosphosites with previously characterized molecular functions and those known to be involved in disease. Although adequate methods were in place to prevent a biased scoring system, it is possible the score may be influenced by social bias to better characterize certain functional sites used for training - such as the tendency for researchers to study conserved positions. While we think such biases are possible, we don’t think they dominate the score as several high scoring sites not included in the training set were supported by additional evidence. The functional score has also shown potential to interpret the impact of mutations in phosphorylated residues. Although we have shown how highly scored phosphorylations are less likely to be mutated or more prone to cause diseases, the full extent of this prioritization hasn’t been explored. Disease variants on functional phosphosites could offer a mechanistic explanation for the disease, possibly indicating a therapeutic strategy perhaps linked to an actionable regulatory kinase.

Together with the full list of human phosphosites and the spectral data released, the phosphosite functional score and the relevant features associated constitute a systematic resource to understand human signaling on a genomic scale. We have shown examples of how this information can prime the design of experiments in order to identify novel regulatory phosphosites. We have shown that high scoring phosphosites might also govern relevant biological processes, as in the case of neuronal differentiation by SMARCC2 phosphorylation. While some of the features may suggest a possible regulatory mechanism, such as the presence of phosphosites at interface positions, the phosphosite functional score itself does not predict mechanisms. Moreover, the phosphosite functional score might not accurately prioritize regulatory sites acting in groups, despite including features that capture local regulatory effects. Additional experimental and computational approaches need to be developed to study the biochemical consequences of individual and coordinated phosphorylations on a large scale.

Not all residue phosphorylations are likely to contribute equally to organismal fitness much the same way that not all transcription factor binding sites, transcripts or even coding genes contribute equally to fitness. Some have speculated that a fraction of phosphosites may serve no purpose at all in present-day species but exist due to the fast-evolutionary changes observed for protein phosphorylation (Landry, Levy, and Michnick 2009). Regardless of the degree of non-functional phosphorylation that may exist it is clear that there is a bottleneck in the functional characterization of phosphosites. The human phosphoproteome and functional score prioritization provided here provide a roadmap to study proteins involved in almost any cellular process.

## Acknowledgements

This study would have been impossible without the selfless deposition of data from hundreds of authors. We would like to extend our gratitude to every one of them. We thank Dr. Jurgen Cox for his insightful advice on the site-decoy multiple testing correction. We would like to thank the members of the Beltrao group for their support collecting features and their relevant comments. This study has been funded by EMBL core funding, and by the Wellcome Trust [grants numbers WT101477MA and 208391/Z/17/Z]. P.B. and D.O. are supported by a Starting Grant Award from the European Research Council (ERC-2014-STG 638884 PhosFunc).

## Abbreviations

AUC: Area Under Curve
CI: Confidence Interval
CPTAC: Clinical Proteomic Tumor Analysis Consortium
CV: Cross-Validation
ESC: Embryonic Stem Cell
KO: Gene Knockout
ML: Machine Learning
MS: Mass Spectrometry
PCA: Principal Component Analysis
ROC: Receiver Operating Characteristic
S, T and Y: Serine, Threonine and Tyrosine
TCGA: The Cancer Genome Atlas
TF: Transcription Factor
PSP: PhosphoSitePlus
WT: Wild Type

## Methods

### Human phosphoproteome MS search

A list of all 307 human datasets annotated to contain phosphorylations was retrieved from the PRIDE database (June 2017) (Vizcaíno et al. 2016). After extensive manual curation of all experimental designs, only untargeted assays that employed phospho-enrichment strategies (e.g. metal oxide affinity chromatography, anti-P-Tyr antibodies, etc.) were included. Datasets with proteomes from more than one species (i.e. infections, xenografts, etc.) were discarded in order to prevent cross-species contaminants. Similarly, datasets with major genetic modifications were also discarded for further analysis. For the remaining 110 PRIDE datasets, additional manual curation was required in order to annotate the biological origin and the search parameters for each raw file. After applying all filters, 6,801 MS raw files remained, corresponding to an accumulated instrument time of 575 days.

All 6,801 MS raw files were jointly searched using MaxQuant 1.6.0.13 (MQ) (Cox and Mann 2008) Andromeda engine (Cox et al. 2011) against the UniProt Human Reference Proteome (71,567 sequences, accessed May 2017) (UniProt Consortium 2018). These were also supplemented with common laboratory contaminants provided by MQ. Based on the labeling strategy or the digestion enzymes, we identified 17 different search parameter groups that were applied to their corresponding raw files. MQ default search parameters as described next include 1% PSM FDR, minimum Andromeda score of 40 and minimum delta score of 6 for modified peptides. Cysteine carbamidomethylation was set as a fixed modification, while oxidation of methionine (M), protein N-terminal acetylation, and phosphorylation of serine (S), threonine (T) and tyrosine (Y) as variable modifications. Minimum peptide length was set to 7 amino acids, and peptides were allowed to have a maximum of two missed-cleavages. All default parameters were also applied for the Orbitrap instruments, the precursor mass tolerance was set to 20 ppm for the first search and 4.5 for the main search. Fragment mass tolerances were set to 20 ppm and 0.5 Da for FT and IT detectors respectively. All other mass tolerance settings were as well kept at default values. The search took 60 days in a Dell PowerEdge R730 rack server configured with Intel(R) Xeon(R) CPU E5-2697 v4 @ 2.30GHz, 64 cores, 256 GB high-capacity DDR4 memory, and 20TB fast access storage drive.

In order to further characterize the false discovery correction at the site level, two separate searches were performed one keeping the default 1% site decoy correction and another search removing this filtering. Despite both searches were corrected at a 1% PSM FDR, the 1% site-FDR search identified 119,809 phosphorylated STYs (121,896 when including the 0.98% of decoys), whereas the uncorrected search identified 252,189 including the 18.48% of decoy hits. This difference highlights the importance of a proper multiple testing correction in all combined searches.

All processing settings and results, including raw and MQ intermediate files, are available in PRIDE under the accession PXD012174.

### Functional annotation of the human phosphoproteome

#### Phosphorylation site MS supporting evidence

A collection of MS-derived phosphosite attributes with the potential to discriminate functional sites was collected for every phosphosite and integrated into the functional score. Regarding the quality of the MS identification within the PRIDE search, we included the best Posterior Empirical Probability (PEP) and the best localization probability from all MS/MS identifications supporting the site with at least a localization probability of 0.5. Separately, a set features were included intending to capture the recurrence of the phosphorylation. These include the total number of FDR-corrected supporting spectral counts with a localization probability greater than 0.5, the number of PRIDE datasets where the site is identified and the number of different cell lines or tissues according to the classification illustrated in Figure 1. Another feature, the number of neighboring PTMs (+/-10 residues) included in PhosphoSitePlus, provides information about the tolerance for modifications and the selective pressure in the protein region (Reimand, Wagih, and Bader 2015). Because of the dependence of some of these features with the number of copies of the protein expressed, we also included in the predictor the MS-derived human consensus protein abundance deposited in PaxDb 4.1 (Wang et al. 2015). Finally, the canonical protein length was as well integrated as reported by UniProt and included in the PRIDE MS search.

#### Residue conservation

As a proxy for residue conservation, we integrated into the functional score for the acceptor residue SIFT score (Vaser et al. 2016). This score predicts the functional impact of missense variants based on sequence homology and the physicochemical properties of the amino acids. The score, ranging from 0 to 1, predicts deleterious variants when scored below 0.05. Since, the conservation of the phosphoacceptor residues does sometimes not only restrict to the acceptor residue, but also to the flanking residues recognized by kinases, we included in the functional score the Evolutionary statistical energy (ΔE) from EVmutation (Hopf et al. 2017), a recent methodology that includes epistatic effects between positions when inferring the tolerance to missense variants. Both SIFT and EVmutation scored the effect of variants instead of providing a single score per residue. In order to collapse the information into a single value per acceptor residue, we used different strategies: the score of the alanine mutant, the average score of mutating to a negatively charged amino acid (E/D), the minimum score and the average score of all variants. Only the score for the alanine variants was included in the final model, as it performed as the best independent predictor of phosphosite function.

#### Human phosphosite age reconstruction

In this study, we applied for the first time a method to reconstruct the ancestral age of human phosphosites. The method originally applied to fungal species (Studer et al. 2016), combines phylogenetic information with cross-species phosphoproteomics in order to infer the most likely ancestor of a given phosphorylation site. To this end, we compiled a list of publicly available MS-identified phosphorylation sites in human (aforementioned PRIDE MS search), *Mus musculus* (108,640 sites), *Rattus norvegicus* (4,605), *Danio rerio* (3,870), *Drosophila melanogaster* (65,480), *Caenorhabditis elegans* (5,544) and *Saccharomyces cerevisiae* (15,708). All sites in every species were re-mapped from their original searches to the same reference proteome by only allowing exact matches of the MS-identified peptide. To ease the inference of more granular distances, also intermediate species were considered despite the lack of available phosphoproteomic data (i.e. *Equus caballus, Gallus gallus* and *Xenopus tropicalis*). Protein sequences, alignments, and phylogenetic trees were all downloaded from Ensembl Compara v86 (Zerbino et al. 2018; Herrero et al. 2016). Next, every phospho-acceptor residue in every species was scored based on its potential to bind a human known kinase-specificity motif. This score was calculated by training a support vector machine (SVM) using as features the sequence-based affinity to 51 Serine/Threonine kinase models integrated into NetPhorest (Horn et al. 2014). The SVM was trained using SVM^ligth^ 6.02 (Joachims 2006) on a balanced set of 4,000 random known phosphosites as positives and the same number of phospho-acceptor residues not know to be phosphorylated as negatives. Since the focus was to reconstruct the age of the phosphosites from a human signaling perspective, all phosphorylated Serines and Threonines were scored using the human ST model, in contrast to the original method that used a different model for each species. Once the phosphorylation potential is estimated from the SVM, the ancestral reconstruction is estimated for every human phosphosite using a maximum likelihood (ML) inference for continuous variables implemented in the R package APE v5.1 (Paradis and Schliep 2018). Intuitively, every residue in the alignment column where a human phosphosite lies is scored: 1.0, if phosphorylation is determined by MS; a value between 0.0 and 1.0 based on the SVM-calculated phosphorylation potential, 0.5 for tyrosines and 0.0 for non-STY residues. After applying ML, we considered as phosphorylated every ancestral node in the phylogenetic tree with a probability score equal or higher than 0.46. Similar to the original methodology, we estimated the phosphosite age based only in the alignment column of the human phosphosite or, alternatively, based on a more flexible region of +/- 3 residues. These 2 metrics were used separately to feed the functional score predictor. Species divergence times were obtained for each ancestral origin from TimeTree.org (Kumar et al. 2017). The pipeline to reproduce the inference of the human phosphosite age is available online (https://github.com/evocellnet/ptmAge).

#### Consensus motif binding

Another phosphosite feature included in the functional score quantifies the similarity of the acceptor residue flanking region to the known kinase-substrate recognition motifs. To first define the kinase specificities, Position Weight Matrices (PWM) were calculated using a +/-7 residue window and equiprobable relative priors for the 143 kinases with at least 10 known human substrates in the PhosphoSitePlus database (Dec 16) (Hornbeck et al. 2019). A matrix similarity score (MSS) ranging from 0 to 1 was next computed between the PWMs and each Serine, Threonine, and Tyrosine in the human proteome as described in the MATCH algorithm (Kel et al. 2003). To collapse all this information into a score for every acceptor residue, we benchmarked the functional phosphorylation predictive power of the highest MSS score from all kinases and, separately, the number of matching kinases where the site scores within the top 0.5%, 1% and 2% of all potential substrates. The highest MSS displayed the best separation between functional and non-functional sites and was the only one included in the final model.

In parallel, we also used a similar strategy to score every acceptor residue based on the posterior probability that a site is recognized by the bundled consensus set of motifs integrated into NetPhorest 2.1 (Horn et al. 2014). Netphorest not only includes kinase recognition motifs but also conserved phosphorylation-dependent binding domains. For every acceptor residue, the highest posterior probability of all models and only kinase models were used as separate features and included in the model. Similarly, we also included as a binary feature whether the flanking region of every acceptor residue matches a Eukaryotic Linear Motif (ELM) (Dinkel et al. 2016).

#### Conditional phospho-regulation

Catalytic regulation of protein phosphorylation has proved to discriminate phosphosite regulatory functions. Based on a previous collection of 41 quantitative human phosphoproteomics studies containing 435 perturbations (Ochoa et al. 2016), we annotated every phosphorylated residue based on the number of conditions in which the site displayed differential phosphoregulation. In order to define the sets of regulated sites based on different levels of stringency, we evaluated separately the number of conditions in which the site is within the top 1%, 5% or 10% of regulation. To avoid correlated features, the 5% cutoff was the only one included in the model as it displayed the best separation for functional phosphosites.

Alternatively, we applied the previously reported inference of kinase activities based on phosphoproteomics data (Hernandez-Armenta et al. 2017), to find co-regulation between the quantified phosphosites and the changes in the activity of 215 human kinases. We included as feature a boolean vector whether the correlation significance after Bonferroni correction is below 0.05. This vector overperformed other explored options to discriminate functional sites, such as the most significant kinase p-value or the number of significantly co-regulated kinases.

#### One-dimensional structural properties

Several structural properties that can be estimated from the protein sequence and summarized as an individual structural descriptor for every acceptor residue were included in the functional score. Using Disopred v2 (Jones and Cozzetto 2015), we estimated the probability that every acceptor residue is natively disordered. The disorder probabilities and a binary classification based on a 0.5 probability cutoff were integrated as features. Solvent accessibility, buried or exposed, inferred from protein sequence was also included, as predicted by the version of ACCpro (Pollastri, Baldi, et al. 2002) implemented in the SCRATCH suite (Cheng et al. 2005). Finally, secondary structure was also estimated from sequence using the neural network-based three-class classification (helix, strand and the rest) contained in SSpro and the full DSSP 8-class output classification implemented in SSpro8 (alpha-helix, 3-10 helix, pi-helix, extended strand, beta-bridge, turn, bend and the rest) (Pollastri, Przybylski, et al. 2002). Due to problems in the memory allocation of the dependent legacy psi-blast software, these features were not estimated for residues in Titin and Mucin-16.

#### Phosphorylation structural hotspots

Recurrence of phosphorylation within the same structural region has been reported as indicative of function (Beltrao et al. 2012). To define these regions, a previous study comprehensively mapped 34,261 phosphorylation sites across 40 eukaryotic species into well-characterized protein domain families (Strumillo et al. 2018). By integrating these data, phosphorylation hotspots were inferred as conserved domain regions with significant enrichment in phosphorylation over random expectation (Implementation available in https://github.com/evocellnet/ptm_hotspots). Here, we integrated as an extra feature in the model, the presence or not of the phospho-acceptor in a hotspot region requiring a p-value < 3.36e-07 and at least 10 MS-identified phosphorylations in the region flanking region (+/- 2 residues). Although this method denoted good discriminative power for functional sites, for the purpose of the functional score, it’s only limited to the 78,390 STYs (5% of all STYs) falling in the well-defined domains. For benchmarking purposes, the remaining STYs were considered as not in a hotspot.

#### Effect on protein stability and interaction interfaces

The predicted protein stability tolerance for genetic variation of the acceptor residue was calculated following a previously described pipeline (Wagih et al. 2018). Structural data were collected from the best human experimentally resolved structures contained in the Protein Data Bank (wwPDB consortium 2019) or, alternatively, from the best homology model derived from ModPipe version 2.2.0 (Pieper et al. 2014). Structural coordinates were remapped to UniProt positions using the SIFTS database (Dana et al. 2019). For the 8,870 proteins with structural data, the impact of all possible variants was computed using FoldX v 4.0 (Guerois, Nielsen, and Serrano 2002). An average ΔG for all runs until convergence is calculated and the ΔΔG is computed as the difference between the wild type and the mutant. Different strategies were benchmarked to collapse the variant effect predictions to a unique ΔΔG for every phospho-acceptor: alanine variant effect, mean of acidic residue variants, mean of all variants and maximum of all variants. Resulting ΔΔGs were discretized according to a previous scale based on its impact on stability (Studer et al. 2014). Only the effect of alanine variants and acidic variants displayed enough discrimination to be included in the functional score model.

To define the boundaries of protein-protein interaction interfaces, structural data on experimentally resolved or modeled interactions were downloaded from Interactome3d (Mosca, Céol, and Aloy 2013). Relative solvent accessibility (RSA) was then calculated using NACCESS (Lee and Richards 1971) for every atom in the interaction structure as well as for both partners separately. Interface residues in each partner and interaction model were defined as all residues changing their relative solvent accessibility between the interacting and non-interacting form. The pipeline to extract protein interfaces from Interactome3d is available online (https://github.com/evocellnet/int3dInterfaces). The presence of the acceptor residue on any interface is included as an extra feature in the functional score model. The impact of variants on the interaction stability was also evaluated using FoldX, but it displayed not enough predictive power to be included in the model.

#### Protein topology annotations

A set of curated features informing about a variable size region around the acceptor residue were integrated into the predictor from the UniProt database (UniProt Consortium 2018). These include whether the phospho-acceptor is in a protein domain, specifically in a kinase domain, in the cytosolic region of a transmembrane protein, in a compositionally biased region, in a zinc finger, in a repeated sequence motif or in any other curated motif.

### AI phosphosite functional prioritization

An MS-derived reference phosphoproteome was defined based on the 119,809 FDR-corrected phosphosites identified in the UniProt reference proteome (UniProt Consortium 2018). In order to prevent problems with redundant sequences, we focused our prioritization strategy in the 116,258 sites contained in the subset of 21,009 reviewed proteins within the reference proteome. Every site was annotated based on a comprehensive list containing 85 of the aforementioned features (**Table S2**). The features were filtered as previously described in order to remove correlated features. Continuous variables were centered, scaled, normalized using the Yeo-Johnson power transformation and near zero-variance variables excluded using the R package caret. Only complete features were considered, except for SIFT were a small set of sites from very large proteins required imputation by nearest neighbors. The discrete variables included in the remaining 38 features, were transformed into dummy variables, expanding the list to the final 59 features.

In order to train a machine learning model capable of regressing the functional potential of all phosphosites, we required the phosphoproteome annotated with functional features and a gold standard set of functional phosphosites. From the MS-identified phosphosites, we defined a gold standard using the 2,638 sites with annotated function in the PhosphoSitePlus database (Nov 2017) (Hornbeck et al. 2019) using the remaining sites of unknown function as negatives. A set of regression models were benchmarked using the same strategy. These include random forest, gradient boosting machine, generalized linear models, regularized linear models, Multivariate Adaptive Regression Splines (MARS), and regression trees. The corresponding R packages for each of the models are described in the caret R wrapper. In all cases, a separate predictor for ST and Y residues was built. Nested repeated cross-validation (CV) was used to estimate the generalization error of the underlying model and its (hyper)parameter search. In the inside loop, a 5-fold CV was performed 10 times in order to tune the parameters. In the outside loop, a 3-fold CV was repeated 5 times to quantify the performance of the model using ROC analysis. An extensive search grid was defined for every training/validation/testing cycle and parallelized in a high-performance computing environment.

Despite the small differences in the performance of the best methods, gradient boosting machine was selected as the final model for the functional score, in part due to its more harmonic distribution of scores. The parameters displaying the best results were: 500 trees, interaction depth of 9, 10 minimum observations in a node and shrinkage of 0.0405 for STs and 0.0105 for Tyrosines. Phosphosite functional scores were derived using these parameters as the median values of all scores obtained by 3-fold CV repeated 30 times.

### Phospho-deficient RANBP1 pull-down assay

A 3xFLAG tagged fusion of both, the WT (UniProt P43487 canonical isoform) and phosphomimetic S60E mutant of RANBP1, were synthesized (GENEWIZ). Lentiviral transduction was used to individually introduce these sequences into HEK293T cells, and RANBP1 expression was controlled under a doxycycline-inducible promoter. Cells from 3 biological replicates of each WT and S60E expression cells were harvested. Additionally, 2 biological replicates of control cells not expressing 3xFLAG RANBP1 were harvested, in which no doxycycline was added to the cells. Protein complexes were purified as previously described (Jäger et al. 2011), and analyzed on a Thermo Q-Exactive Plus mass spectrometer. Raw data was searched using MaxQuant (Cox and Mann 2008), and high confidence protein-protein interactions were scored using SAINTexpress (Teo et al. 2014).

### TF activity inference and phosphosite coregulation

To systematically estimate transcription factor activities and phosphosite co-regulation across a panel of primary tumors, paired basal gene expression and quantitative phosphoproteomic data from the same breast tumors were retrieved from The Cancer Genome Atlas (TCGA) (Cancer Genome Atlas Network 2012) and Clinical Proteomic Tumor Analysis Consortium (CPTAC) (Mertins et al. 2016), respectively. From the 105 breast tumors with matched phosphoproteomics data, raw counts were downloaded from the Gene Expression Omnibus (GSE62944) (Edgar, Domrachev, and Lash 2002), normalized, processed into counts per million reads and z-transformed. To infer transcription factor (TF) activities from the expression data, 289 good quality TF regulons (A-C) were retrieved from the OmniPath resource (Türei, Korcsmáros, and Saez-Rodriguez 2016) and an enrichment test per tumor was calculated on them using the analytic Rank-based Enrichment Analysis (aREA) method implemented in Viper (Alvarez et al. 2016). The Normalized Enrichment Score (NES) was used as an estimate of relative TF activity as described in previous studies (Garcia-Alonso et al. 2018). The changes in phosphorylation for TF sites quantified across the 77 samples passing the CPTAC quality control criteria were correlated with their corresponding TF activities.

### Effects of genetic variation in human phosphosites

Variation in natural human populations for 60,706 unrelated individuals was retrieved from the Exome Aggregation Consortium (ExAC) (Lek et al. 2016). The variants with their corresponding adjusted allele frequencies were aligned to UniProt positions using the Needleman-Wunsch global alignment implemented in the Biostrings R package. Only variants observed in at least 10 counts were considered for further analysis. In cases were multiple alleles mapped to the same exact amino acid substitution, the variant with the highest adjusted frequency was preserved. For comparison purposes, Minor Allele Frequency (MAF) for phospho-acceptor residues was calculated as the frequency at which the second most common allele occurs in the population. To further understand the effects in human variation, a total of 159,633 human clinically relevant variants were retrieved from ClinVar (Landrum et al. 2018), 1,784 of which match phospho-acceptor residues. The annotated clinical significances were collapsed into benign, uncertain or pathogenic. Finally, the phenotypic consequences of introducing 22,090 missense mutations in 3,022 proteins were retrieved from UniProt (UniProt Consortium 2018). The 4,764 point-mutations on phospho-acceptor residues were classified into Alanine (A), Phosphomimetic (E or D) or Other depending on the resulting residue. Since the phenotypic consequences are encoded as free text, an in-house parser was required to annotate the effects as gain-of-function, loss-of-function or no effect.

### Phenotypic effects of distant phospho-deficient variants

#### TDH3 phosphomutant construction

Phospho-deficient mutants TDH3 S149A and T151A were constructed by introducing the point mutations into the TDH3 endogenous locus in the Y8205 background strain (MATalpha, his3 1; leu2 0; ura3 0; MET15+; LYS2+; can1::STE2pr-SpHIS5; lyp1::STE3pr-LEU2 + GALpr-SceI-NAT) followed by a URA3 marker after the stop codon. The URA3 marker was flanked by SceI recognition sites and removed by induction of the endonuclease as previously described (Khmelinskii et al. 2011). Point mutations were verified by DNA sequencing.

#### TDH3 Growth curves description

Yeast strains including wild-type, gene control, TDH3 KO from the yeast gene knockout library (Winzeler et al. 1999) and the 2 phospho-deficient mutants were inoculated into a final volume of 100 μL SC media with and without 75µM doxorubicin in 96-well plates (initial OD_660nm_=0.05). 4 biological replicates were inoculated in each plate and the experiment was performed 3 times. All plates were sealed with breathable membranes (Breathe-Easy) and incubated at 30°C in a thermostated incubator (Cytomat 2, Thermo Scientific) with continuous shaking. OD_660nm_ was measured every 30 min for 48h in a Filtermax F5 multimode plate reader (Molecular Devices). Growth curves were estimated by fitting a standard logistic equation included in the Growthcurver R package. Cellular fitness was defined by the area under the curve (AUC).

#### TDH3 Mutant activity measurement

Yeast strains growing in exponential phase were diluted to the same OD (OD_660nm_=0.2) in 3 biological replicates and 1ml was collected by centrifugation. Cell lysis was adapted for yeast cells: cell pellets were resuspended in 500µl of cold lysis buffer (20 mM Tris pH8, 15mM EDTA pH8, 15mM EGTA pH8 and 0.1% Triton X-100). Glass beads were added in equal volume (500µl) and cells were lysed by vortexing at 4°C. Enzymatic activity was quantified twice using the colorimetric Glyceraldehyde-3-Phosphate Dehydrogenase Activity Assay Kit (Abcam) as described by the manufacturer.

### Phenotypicassayofphospho-deficientSMARCC2duringneuronal differentiation

#### Generation of Smarcc2/BAF170 mutant cell lines

Mouse embryonic stem (ES) cells (129XC57BL/6J) were cultured in media containing Knockout-DMEM (Thermo Fisher) with 15% EmbryoMax FBS (Millipore) and 20 ng/ml leukemia inhibitory factor (LIF, produced by Protein Expression Facility at EMBL Heidelberg), 1% non-essential amino acids, 1% Glutamax, 1% Pen/Strep, 1% of 55mM beta-Mercaptoethanol solution. Cells were maintained at 37°C with 5% CO2. ESCs were routinely tested for mycoplasma absence by PCR.

ES lines with double C to G point mutations encoding S302A and S304A substitutions in Smarcc2/BAF170 were generated using CRISPR-Cas9 mediated knock-in mutation according to the published protocol (Ran et al. 2013). Briefly, oligonucleotides encoding guide sequences were cloned into pSpCas9(BB)-2A-GFP (PX458, Addgene). The resulting plasmid (2 μg Smarcc2_g1 or g2) and 100 μM ssODN repair templates (180 bp, ssODN_Smarcc2_g1 or g2, IDT ultrameres) were nucleofected into ES cells using 4D-nucleofector (Lonza) for gene editing. Two days after nucleofection, individual transfected cells were plated in 96 well plates by FACS. Cells carrying homozygous or heterozygous introduced mutations in Smarcc2/BAF170 were identified by Sanger sequencing. In addition, cells that retained a wild type Smarcc2/BAF170 sequence despite having gone through the CRISPR mutagenesis process were retained as CRISPR controls.

Smarcc2_g1 AGGGGGGCAACTATAAGAAG

Smarcc2_g2 CTCTGGGGTGGGTGAAGGAG

ssODN_Smarcc2_g1

GGGTAAACTGCTGCTCATCCCACGTCCTGTTGTGCAGGTAAACAGCCCAG

ATTCAGACAGACGAGACAAGAAGGGGGGCAATTATAAGAAAAGAAAGCGC

GCTCCCGCTCCTTCACCCACCCCAGAGGCTAAGAAGAAAAACGCTAAGAA

AGGGTAAGCTACCTCCTGTGCCCACACGCG

ssODN_Smarcc2_g2

TCCCACGTCCTGTTGTGCAGGTAAACAGCCCAGATTCAGACAGACGAGAC

AAGAAGGGGGGCAACTATAAGAAGAGGAAGCGCGCTCCCGCCCCATCACC

CACCCCAGAGGCTAAGAAGAAAAACGCTAAGAAAGGGTAAGCTACCTCCT

GTGCCCACACGCGCTGTCCTGATGCCATCTCA

#### Neuronal Differentiation

ES cells with heterozygous or homozygous Smarcc2 mutations, along with CRISPR control cells, were neuronally differentiated according to Bibel et al. To start differentiation, ES cells were plated on bacterial Petri dishes in CA media containing DMEM high glucose (Thermo Fisher) with 10% FBS, 1% non-essential amino acids, 1% Glutamax, 1% Pen/Strep, 1% of 55mM beta-Mercaptoethanol solution. After 4 days retinoic acid was added to embryonic bodies at a final concentration of 5µM. For neural culture, after 8 days embryoid bodies were dissociated with trypsin and plated in N2 media composed of regular DMEM supplemented with N2 and B27-VitaminA (Thermo Fisher). Plates were pre-coated with Poly-D-Lysine (Sigma) and Laminin (Roche). Half of the N2 media was changed every two days. Samples of differentiating cells were snap frozen in liquid nitrogen on the day of plating (D0) as well as on the four (D4), eight (D8) and twelve (D12) days following differentiation.

#### Western Blotting

ES cell-derived neuronal BAF170 protein levels were visualized by western blot. Crude nuclear extracts were prepared from D12 mutant cell lines and CRISPR controls. Extracts were also prepared from D0, D4, D8 and D12 wild type differentiating cells. Extracts were run on SDS/PAGE gels (4-12%), transferred to PVDF membranes and probed with primary antibodies to BAF170 (Active Motif, 1:1000) and Histone H4 (1:6000, Abcam). Membranes were washed, probed with HRP tagged goat anti-rabbit secondary antibodies, and visualized with Immobilon chemiluminescent reagent (Millipore) using a Chemidoc Touch imaging system (Bio-Rad).

#### Immunofluorescence

Neuronal morphology and SMARCC2/BAF170 protein localization were visualized using indirect immunofluorescence of D12 neuronal cultures. D8 cells were placed on coverslips and fixed on D12 with 2% PFA in PBS for 15 minutes at room temperature. Coverslips were washed 3 times in PBS and permeabilized with 100% methanol for 5 minutes at −20°C. Permeabilized cells were washed twice with wash buffer (0.05% Tween 20 in PBS) and blocked for 30 minutes with 2% BSA in PBS. Cells were then incubated with primary antibodies (1:400 MAP2 (Sigma) and 1:2000 BAF170 (Active Motif)) for 1 hour at room temperature or overnight at 4°C. Coverslips were given 3 washes in wash buffer for 5 minutes each. Secondary antibodies (anti-Mouse Alexa Fluor 594 and anti-Rabbit Alexa Fluor 488) were applied for 30 minutes at room temperature followed by an additional two 5-minute washes in wash buffer. Coverslips were washed briefly in sterile water, inverted and mounted on microscope slides with Prolong Gold antifade reagent. Images were collected at 40x with a Ti-Eclipse Fluorescence Microscope (Nikon). Image Analysis was performed using Fiji and the Cell Counter plugin.

#### RNA sequencing

Total RNA was extracted from D8 and D12 differentiating cells using the RNeasy RNA isolation kit (Qiagen) followed by DNase digestion using TURBO DNase (Thermo Fisher). mRNAs were isolated from 1 μg of total RNA using a PolyA selection kit (NEB) and sequencing libraries were prepared using the NEBnext Ultra RNA library prep kit for Illumina (NEB). Completed libraries were sequenced on a NextSeq 500 sequencer (Illumina) using a 75 bp single-end run. Reads were aligned to the murine GRCm38 assembly using HISAT2 v2.1.0 (Kim, Langmead, and Salzberg 2015). Aligned reads were converted to BAM format and sorted using samtools v0.1.19 (Li et al. 2009). Read counts were calculated using htseq-count (Anders, Pyl, and Huber 2015) using the murine gene sets annotated in Ensembl v93 (Zerbino et al. 2018). Reads with less than 10 counts were discarded for further analysis. Read counts from all samples were combined using the Rsubread (Liao, Smyth, and Shi 2013) and analyzed using DESeq2 (Love, Huber, and Anders 2014). Gene set signatures for ESC, neuronal differentiation and TF regulons were downloaded from MSigDB (Liberzon et al. 2015).

---

**Figure S1.**
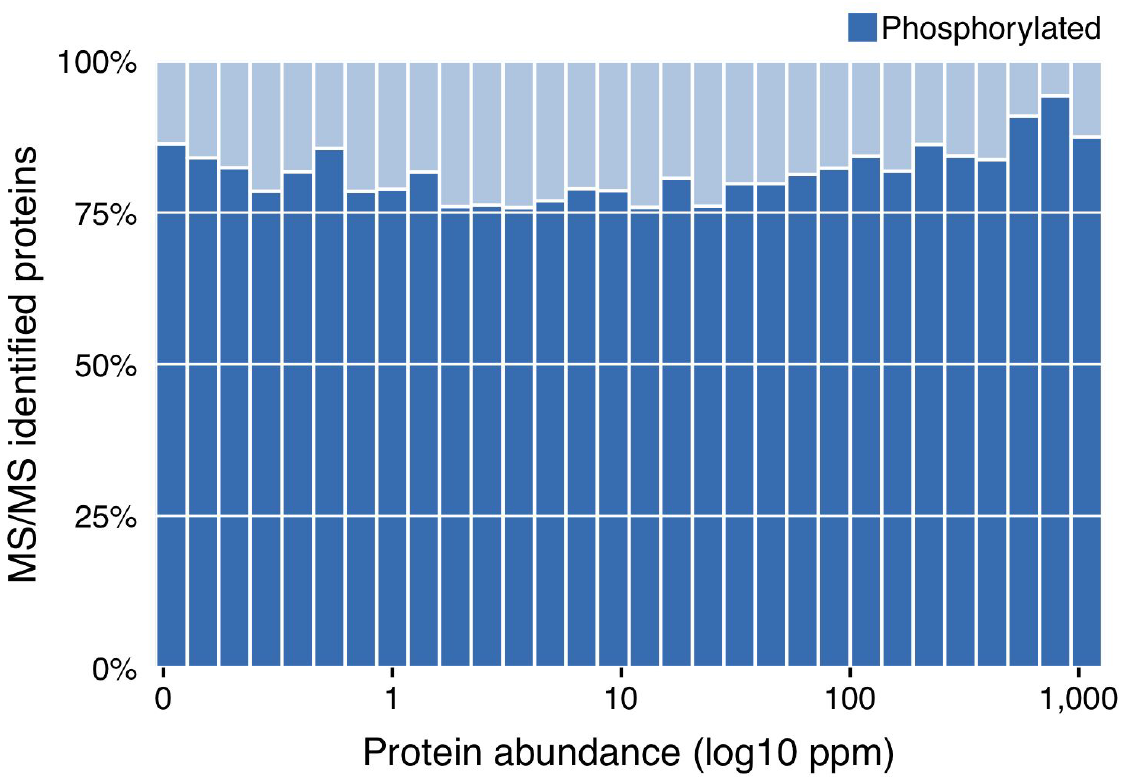
Ratio of phosphorylated proteins binned by the protein consensus abundance. At least one identified phosphorylation was required to consider a protein phosphorylated. Protein abundance data obtained from the PaxDb consensus human proteome.

**Figure S2.**
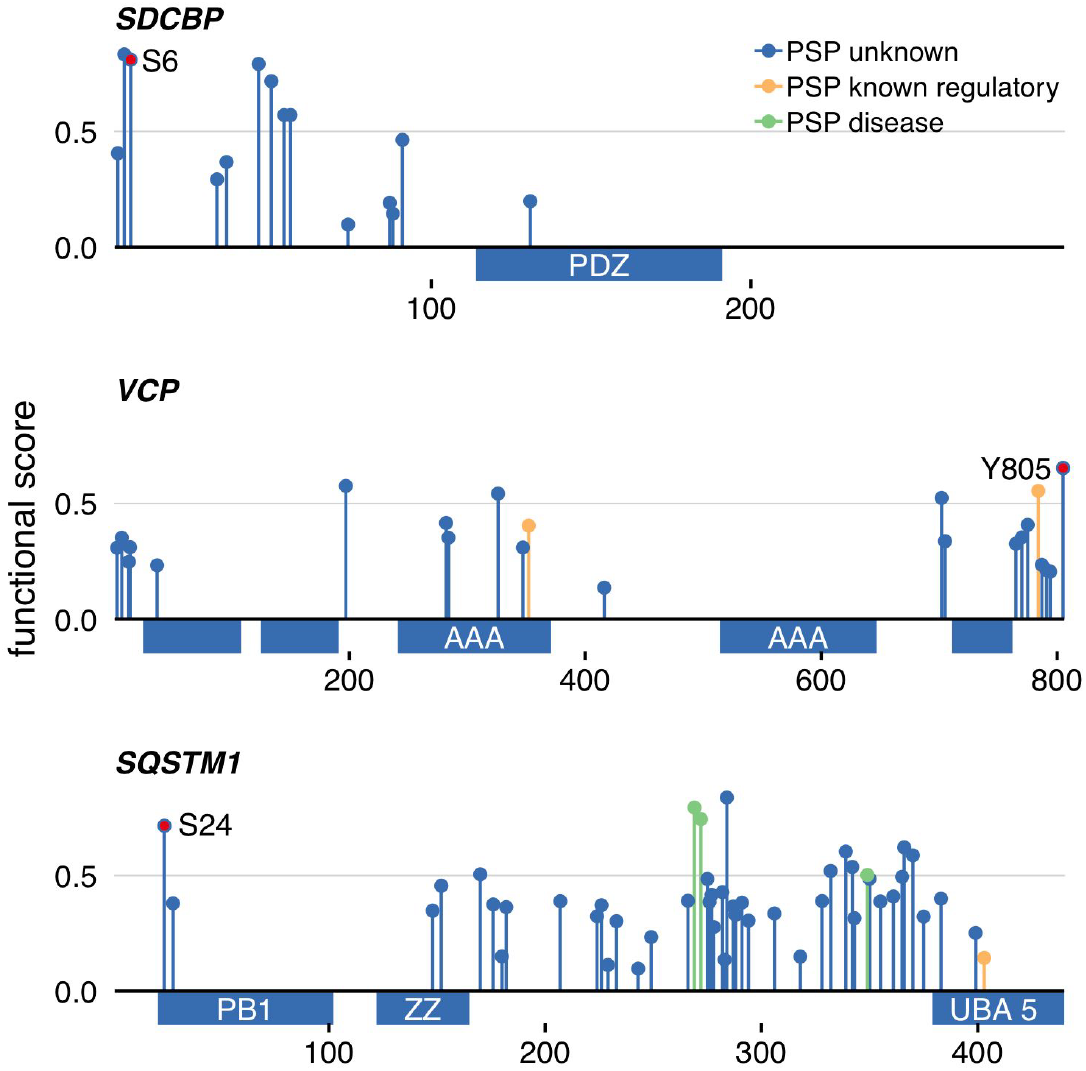
Examples of experimentally validated phosphosites not included in the training set. Functional score and position for phosphosites identified in the PRIDE search and colored by the level of functional annotation in PhosphositePlus (PSP). Sites marked in red represent sites of unknown function in PSP that were supported by experimental evidence in the literature.

**Figure S3.**
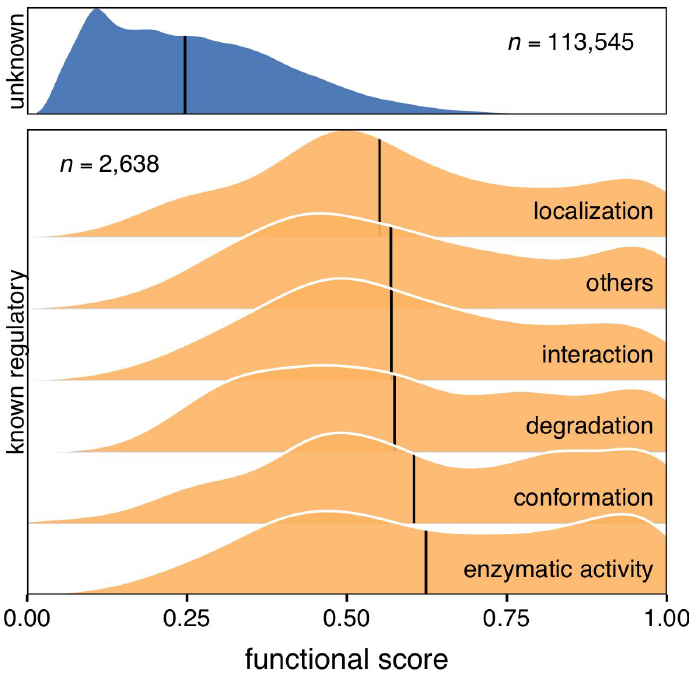
Separation of sites of known and unknown function based on their molecular regulatory role. The molecular function was obtained from PSP.

**Figure S4.**
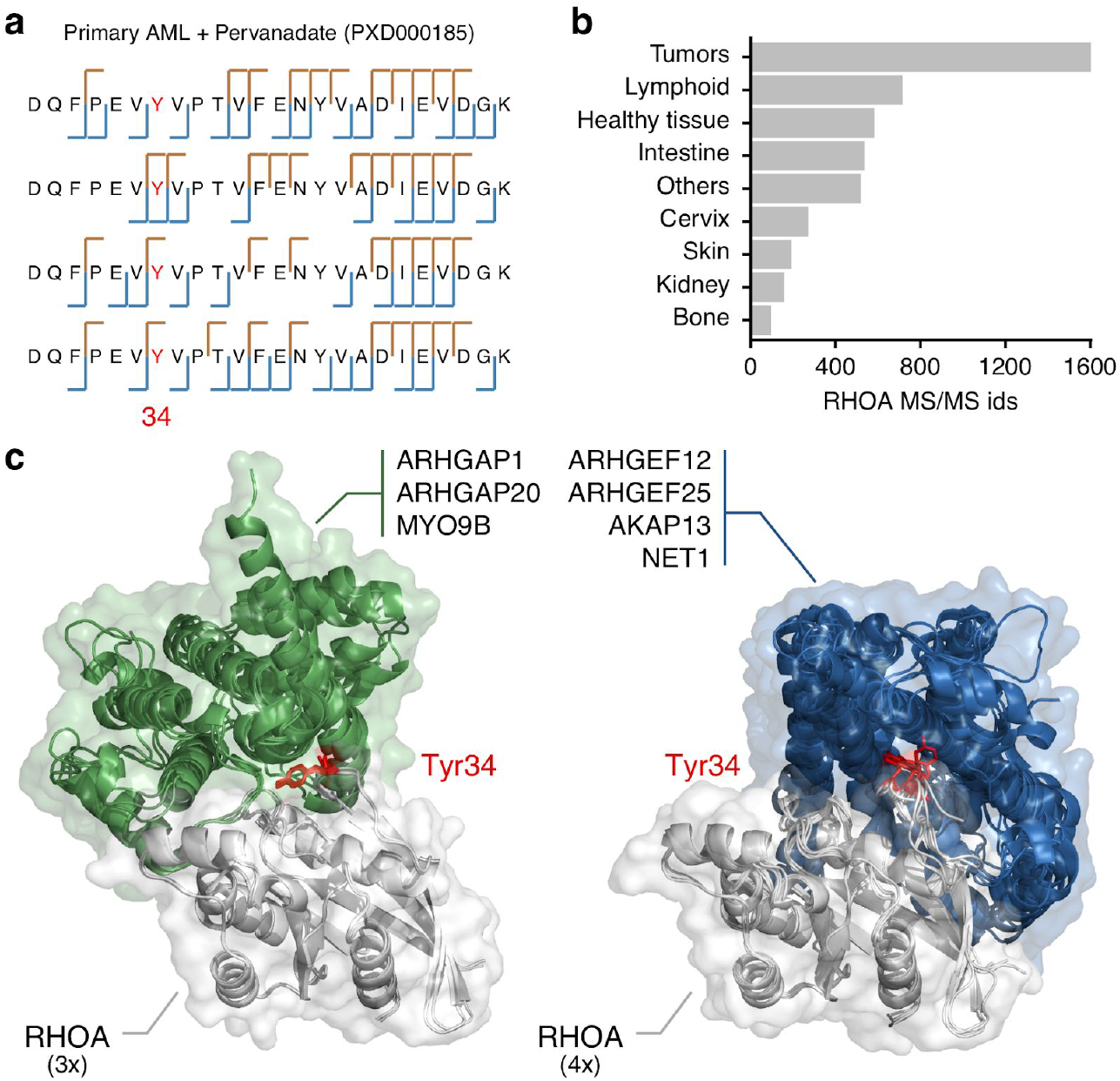
Identification and characterization of known regulatory sites determining protein binding specificity. a) b-ions and y-ions for the 4 only RHOA phosphopeptides phosphorylated in Tyr34. The 4 peptides were identified from Primary AML tumors treated with the protein tyrosine phosphatase pervanadate. b) Number of MS/MS identifications in all samples containing modified or unmodified peptides in RHOA. c) Aligned structural data for binding of RHOA with 7 different partners (PDB files: 1tx4, 5hpy and 3msx - left - 4d0n, 4xh9, 2rgn and1tx4 - right). The position of the acceptor tyrosine - red - changes depending on the group of binding partners.

**Figure S5.**
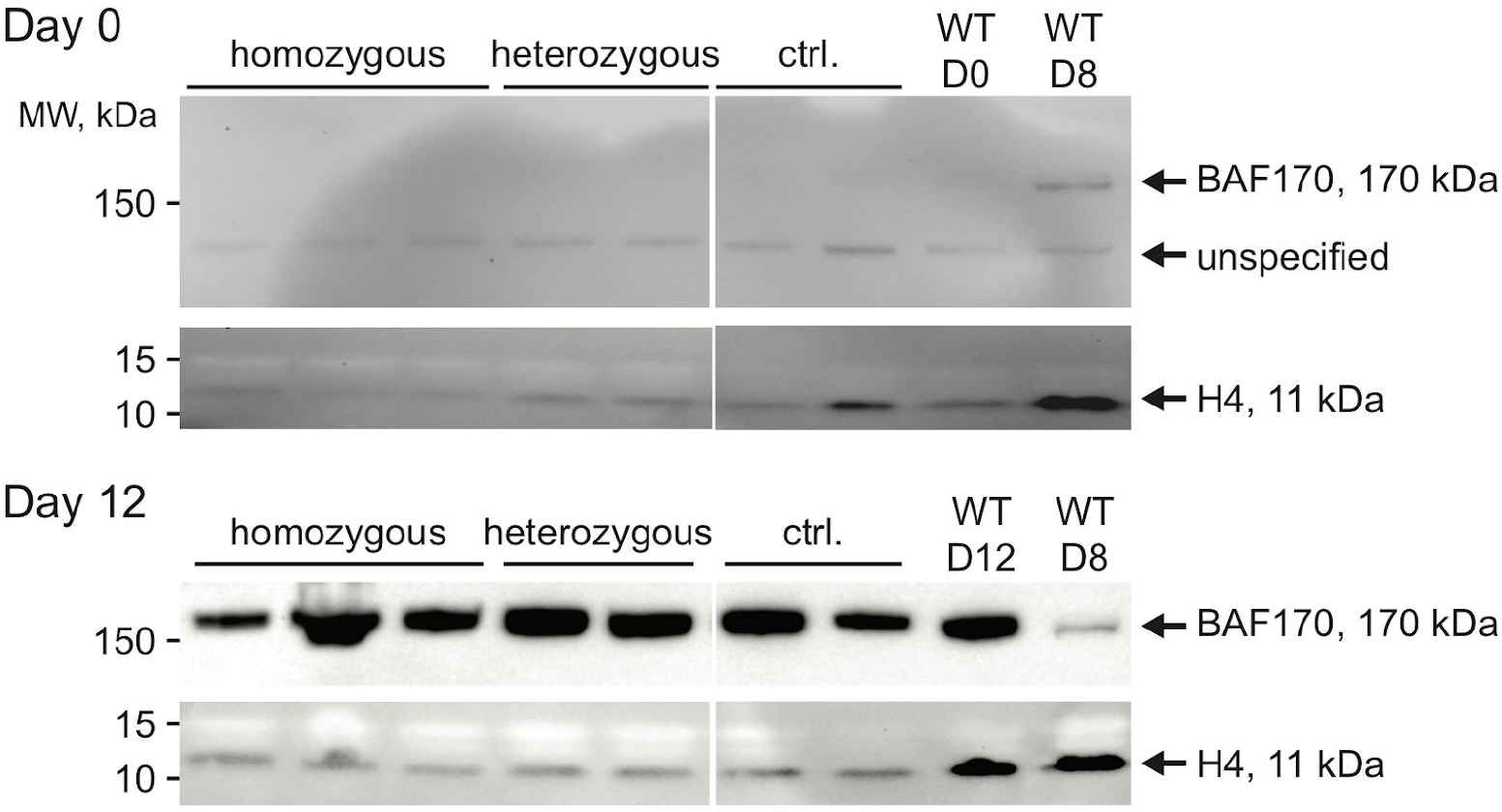
Smarcc2/Baf170 protein is constitutively express day-12 of neuronal differentiation independently of the genetic background.

**Figure S6.**
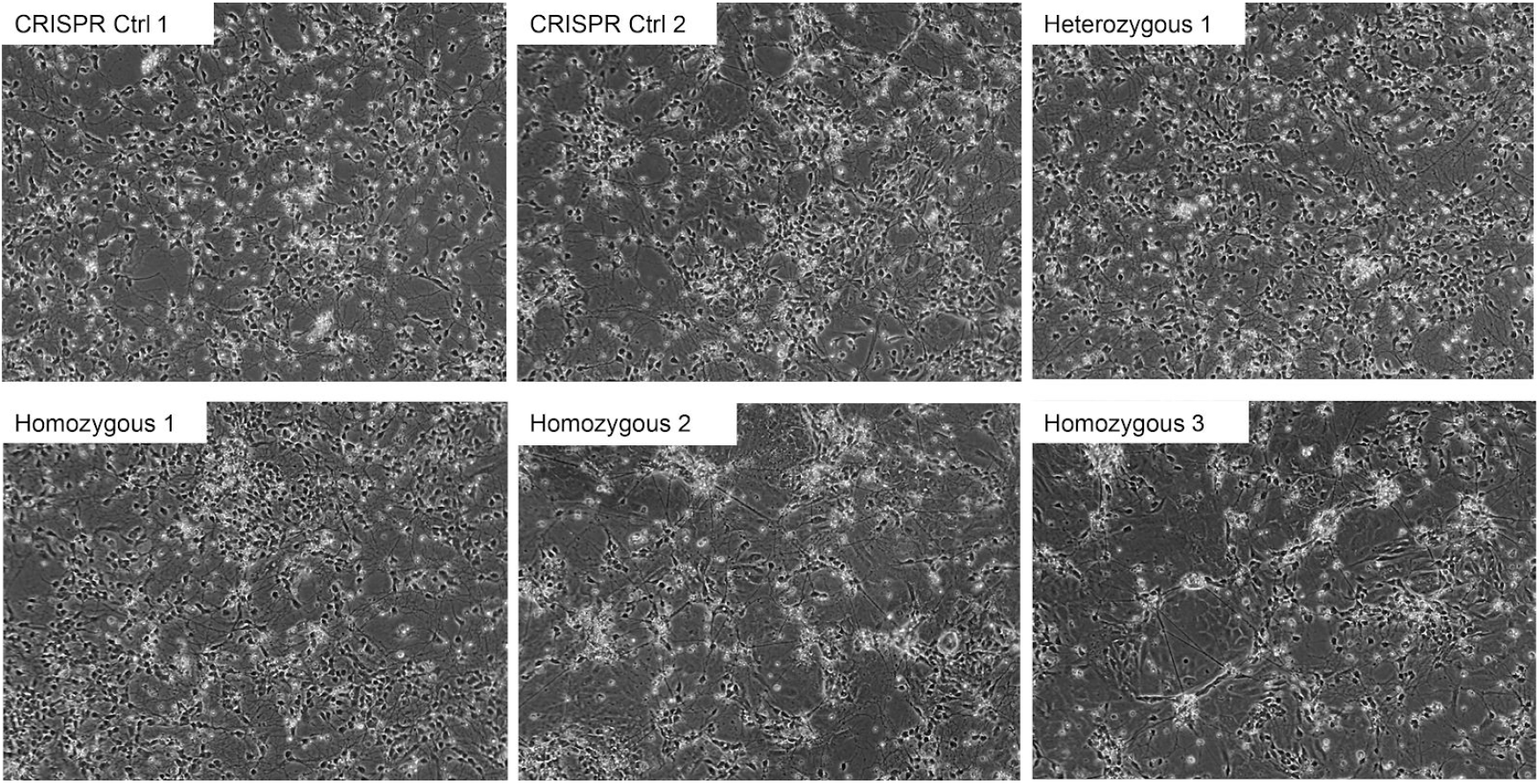
Morphological differences in day-12 neuronal differentiation for Smarcc2 CRISPR control, heterozygous and homozygous S302A/S304A clones. Bright field images, 20x

**Figure S7.**
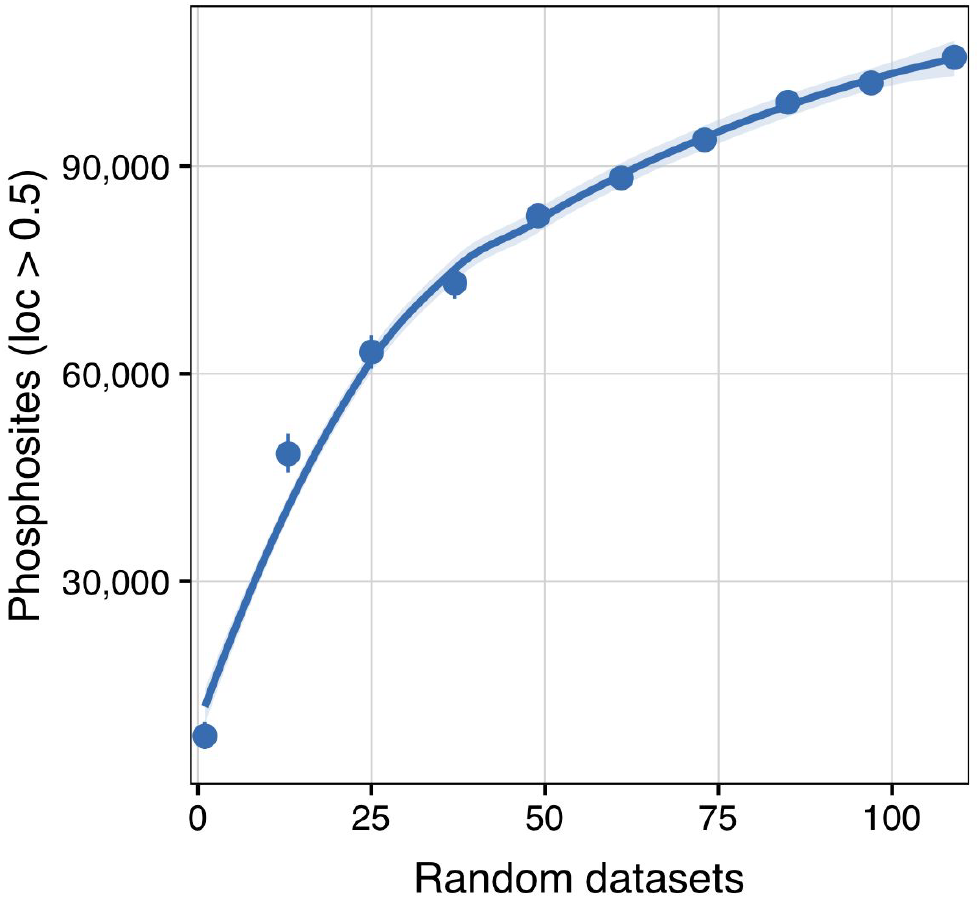
Accumulation of phosphosites as the number of phospho-enriched datasets deposited in PRIDE grows. Rarefaction curve for random samples of datasets. The total number of sites only refer to phosphosites with localization probability greater than 0.5.

## Supplementary Tables

**Table S1. PRIDE data included in the re-analysis**. The spreadsheet includes the PRIDE datasets under study, specifying each of the raw files included in the search and their search parameters.

**Table S2. Annotated phosphoproteome.** Features that might indicate phosphosite function for the 116,258 sites contained in the subset of 21,009 reviewed proteins within the human UniProt reference proteome.

**Table S3. Phosphosite functional scores.** 116,258 scored sites contained in the subset of 21,009 reviewed proteins within the human UniProt reference proteome.

**Table S4. RANBP1 pull down-MS results.**

## References

Aebersold, Ruedi, and Matthias Mann. 2016. “Mass-Spectrometric Exploration of Proteome Structure and Function.” Nature 537 (7620): 347–55.

Alvarez, Mariano J., Yao Shen, Federico M. Giorgi, Alexander Lachmann, B. Belinda Ding, B. Hilda Ye, and Andrea Califano. 2016. “Functional Characterization of Somatic Mutations in Cancer Using Network-Based Inference of Protein Activity.” Nature Genetics 48 (8): 838–47.

Anders, Simon, Paul Theodor Pyl, and Wolfgang Huber. 2015. “HTSeq--a Python Framework to Work with High-Throughput Sequencing Data.” Bioinformatics 31 (2): 166–69.

Beltrao, Pedro, Véronique Albanèse, Lillian R. Kenner, Danielle L. Swaney, Alma Burlingame, Judit Villén, Wendell A. Lim, James S. Fraser, Judith Frydman, and Nevan J. Krogan. 2012. “Systematic Functional Prioritization of Protein Posttranslational Modifications.” Cell 150 (2): 413–25.

Beltrao, Pedro, Peer Bork, Nevan J. Krogan, and Vera van Noort. 2013. “Evolution and Functional Cross-Talk of Protein Post-Translational Modifications.”; Molecular Systems Biology; 9 (December): 714.

Betts, Matthew J., Oliver Wichmann, Mathias Utz, Timon Andre, Evangelia Petsalaki, Pablo Minguez, Luca Parca, et al. 2017. “Systematic Identification of Phosphorylation-Mediated Protein Interaction Switches.” PLoS Computational Biology 13 (3): e1005462.

Cahoy, John D., Ben Emery, Amit Kaushal, Lynette C. Foo, Jennifer L. Zamanian, Karen S. Christopherson, Yi Xing, et al. 2008. “A Transcriptome Database for Astrocytes, Neurons, and Oligodendrocytes: A New Resource for Understanding Brain Development and Function.” The Journal of Neuroscience: The Official Journal of the Society for Neuroscience 28 (1): 264–78.

Cancer Genome Atlas Network. 2012. “Comprehensive Molecular Portraits of Human Breast Tumours. Nature 490 (7418): 61–70.

Cheng, J., A. Z. Randall, M. J. Sweredoski, and P. Baldi. 2005. “SCRATCH: A Protein Structure and Structural Feature Prediction Server.” Nucleic Acids Research 33 (Web Server issue): W72–76.

Christian, Frank, Eberhard Krause, Miles D. Houslay, and George S. Baillie. 2014. “PKA Phosphorylation of p62/SQSTM1 Regulates PB1 Domain Interaction Partner Binding.” Biochimica et Biophysica Acta 1843 (11): 2765–74.

Cox, Jürgen, and Matthias Mann. 2008. “MaxQuant Enables High Peptide Identification Rates, Individualized P.p.b.-Range Mass Accuracies and Proteome-Wide Protein Quantification.” Nature Biotechnology 26 (12): 1367–72.

Cox, Jürgen, Nadin Neuhauser, Annette Michalski, Richard A. Scheltema, Jesper V. Olsen, and Matthias Mann. 2011. “Andromeda: A Peptide Search Engine Integrated into the MaxQuant Environment.” Journal of Proteome Research 10 (4): 1794–1805.

Dana, Jose M., Aleksandras Gutmanas, Nidhi Tyagi, Guoying Qi, Claire O’Donovan, Maria Martin, and Sameer Velankar. 2019. “SIFTS: Updated Structure Integration with Function, Taxonomy and Sequences Resource Allows 40-Fold Increase in Coverage of Structure-Based Annotations for Proteins.” Nucleic Acids Research 47 (D1): D482–89.

Dinkel, Holger, Kim Van Roey, Sushama Michael, Manjeet Kumar, Bora Uyar, Brigitte Altenberg, Vladislava Milchevskaya, et al. 2016. “ELM 2016--Data Update and New Functionality of the Eukaryotic Linear Motif Resource.” Nucleic Acids Research 44 (D1): D294–300.

Edgar, Ron, Michael Domrachev, and Alex E. Lash. 2002. “Gene Expression Omnibus: NCBI Gene Expression and Hybridization Array Data Repository.” Nucleic Acids Research 30 (1): 207–10.

Friedman, Jerome H. 2002. “Stochastic Gradient Boosting.” Computational Statistics & Data Analysis 38 (4): 367–78.

Garcia-Alonso, Luz, Francesco Iorio, Angela Matchan, Nuno Fonseca, Patricia Jaaks, Gareth Peat, Miguel Pignatelli, et al. 2018. “Transcription Factor Activities Enhance Markers of Drug Sensitivity in Cancer.” Cancer Research 78 (3): 769–80.

Guerois, Raphael, Jens Erik Nielsen, and Luis Serrano. 2002. “Predicting Changes in the Stability of Proteins and Protein Complexes: A Study of More than 1000 Mutations.” Journal of Molecular Biology 320 (2): 369–87.

Hailesellasse Sene, Kagnew, Christopher J. Porter, Gareth Palidwor, Carolina Perez-Iratxeta, Enrique M. Muro, Pearl A. Campbell, Michael A. Rudnicki, and Miguel A. Andrade-Navarro. 2007. “Gene Function in Early Mouse Embryonic Stem Cell Differentiation.” BMC Genomics 8 (March): 85.

Hernandez-Armenta, Claudia, David Ochoa, Emanuel Gonçalves, Julio Saez-Rodriguez, and Pedro Beltrao. 2017. “Benchmarking Substrate-Based Kinase Activity Inference Using Phosphoproteomic Data.” Bioinformatics 33 (12): 1845–51.

Herrero, Javier, Matthieu Muffato, Kathryn Beal, Stephen Fitzgerald, Leo Gordon, Miguel Pignatelli, Albert J. Vilella, et al. 2016. “Ensembl Comparative Genomics Resources.” Database: The Journal of Biological Databases and Curation 2016 (February). https://doi.org/10.1093/database/bav096.

Hopf, Thomas A., John B. Ingraham, Frank J. Poelwijk, Charlotta P. I. Schärfe, Michael Springer, Chris Sander, and Debora S. Marks. 2017. “Mutation Effects Predicted from Sequence Co-Variation.” Nature Biotechnology; 35 (2): 128–35.

Hornbeck, Peter V., Jon M. Kornhauser, Vaughan Latham, Beth Murray, Vidhisha Nandhikonda, Alex Nord, Elzbieta Skrzypek, Travis Wheeler, Bin Zhang, and Florian Gnad. 2019. “15 Years of PhosphoSitePlus(r): Integrating Post-Translationally Modified Sites, Disease Variants and Isoforms.” Nucleic Acids Research; 47 (D1): D433–41.

Horn, Heiko, Erwin M. Schoof, Jinho Kim, Xavier Robin, Martin L. Miller, Francesca Diella, Anita Palma, Gianni Cesareni, Lars Juhl Jensen, and Rune Linding. 2014. “KinomeXplorer: An Integrated Platform for Kinome Biology Studies.” Nature Methods 11 (6): 603–4.

Houssa, B., J. de Widt, O. Kranenburg, W. H. Moolenaar, and W. J. van Blitterswijk. 1999. “Diacylglycerol Kinase Theta Binds to and Is Negatively Regulated by Active RhoA.” The Journal of Biological Chemistry 274 (11): 6820–22.

Hwang, Hyo-In, Jae-Hoon Ji, and Young-Joo Jang. 2011. “Phosphorylation of Ran-Binding Protein-1 by Polo-like Kinase-1 Is Required for Interaction with Ran and Early Mitotic Progression.” The Journal of Biological Chemistry 286 (38): 33012–20.

Jäger, Stefanie, Peter Cimermancic, Natali Gulbahce, Jeffrey R. Johnson, Kathryn E. McGovern, Starlynn C. Clarke, Michael Shales, et al. 2011. “Global Landscape of HIV-Human Protein Complexes.” Nature 481 (7381): 365–70.

Jaglin, Xavier Hubert, Karine Poirier, Yoann Saillour, Emmanuelle Buhler, Guoling Tian, Nadia Bahi-Buisson, Catherine Fallet-Bianco, et al. 2009. “Mutations in the Beta-Tubulin Gene TUBB2B Result in Asymmetrical Polymicrogyria.” Nature Genetics 41 (6): 746–52.

Joachims, Thorsten. 2006. Making Large-Scale SVM Learning Practical.

Jones, David T., and Domenico Cozzetto. 2015. “DISOPRED3: Precise Disordered Region Predictions with Annotated Protein-Binding Activity.” Bioinformatics 31 (6): 857–63.

Kanshin, Evgeny, Louis-Philippe Bergeron-Sandoval, S. Sinan Isik, Pierre Thibault, and Stephen W. Michnick. 2015. “A Cell-Signaling Network Temporally Resolves Specific versus Promiscuous Phosphorylation.” Cell Reports 10 (7): 1202–14.

Kel, A. E., E. Gössling, I. Reuter, E. Cheremushkin, O. V. Kel-Margoulis, and E. Wingender. 2003. ”MATCH: A Tool for Searching Transcription Factor Binding Sites in DNA Sequences.” Nucleic Acids Research 31 (13): 3576–79.

Khmelinskii, Anton, Matthias Meurer, Nurlanbek Duishoev, Nicolas Delhomme, and Michael Knop. 2011. “Seamless Gene Tagging by Endonuclease-Driven Homologous Recombination.” PloS One 6 (8): e23794.

Kim, Daehwan, Ben Langmead, and Steven L. Salzberg. 2015. “HISAT: A Fast Spliced Aligner with Low Memory Requirements.” Nature Methods 12 (4): 357–60.

Kumar, Sudhir, Glen Stecher, Michael Suleski, and S. Blair Hedges. 2017. “TimeTree: A Resource for Timelines, Timetrees, and Divergence Times.” Molecular Biology and Evolution 34 (7): 1812–19.

Lahiry, Piya, Ali Torkamani, Nicholas J. Schork, and Robert A. Hegele. 2010. “Kinase Mutations in Human Disease: Interpreting Genotype-Phenotype Relationships.” Nature Reviews. Genetics 11 (1): 60–74.

Landrum, Melissa J., Jennifer M. Lee, Mark Benson, Garth R. Brown, Chen Chao, Shanmuga Chitipiralla, Baoshan Gu, et al. 2018. “ClinVar: Improving Access to Variant Interpretations and Supporting Evidence.” Nucleic Acids Research 46 (D1): D1062–67.

Landry, Christian R., Emmanuel D. Levy, and Stephen W. Michnick. 2009. “Weak Functional Constraints on Phosphoproteomes.” Trends in Genetics: TIG 25 (5): 193–97.

Lee, B., and F. M. Richards. 1971. “The Interpretation of Protein Structures: Estimation of Static Accessibility.” Journal of Molecular Biology 55 (3): 379–IN4.

Lek, Monkol, Konrad J. Karczewski, Eric V. Minikel, Kaitlin E. Samocha, Eric Banks, Timothy Fennell, Anne H. O’Donnell-Luria, et al. 2016. “Analysis of Protein-Coding Genetic Variation in 60,706 Humans.” Nature 536 (7616): 285–91.

Liao, Yang, Gordon K. Smyth, and Wei Shi. 2013. “The Subread Aligner: Fast, Accurate and Scalable Read Mapping by Seed-and-Vote.” Nucleic Acids Research 41 (10): e108.

Liberzon, Arthur, Chet Birger, Helga Thorvaldsdóttir, Mahmoud Ghandi, Jill P. Mesirov, and Pablo Tamayo. 2015. “The Molecular Signatures Database (MSigDB) Hallmark Gene Set Collection.” Cell Systems 1 (6): 417–25.

Li, Heng, Bob Handsaker, Alec Wysoker, Tim Fennell, Jue Ruan, Nils Homer, Gabor Marth, Goncalo Abecasis, Richard Durbin, and 1000 Genome Project Data Processing Subgroup. 2009. “The Sequence Alignment/Map Format and SAMtools.” Bioinformatics 25 (16): 2078–79.

Liu, Jeffrey J., Kirti Sharma, Luca Zangrandi, Chongguang Chen, Sean J. Humphrey, Yi-Ting Chiu, Mariana Spetea, Lee-Yuan Liu-Chen, Christoph Schwarzer, and Matthias Mann. 2018. “In Vivo Brain GPCR Signaling Elucidated by Phosphoproteomics.” Science 360 (6395). https://doi.org/10.1126/science.aao4927.

Love, Michael I., Wolfgang Huber, and Simon Anders. 2014. “Moderated Estimation of Fold Change and Dispersion for RNA-Seq Data with DESeq2.” Genome Biology 15 (12): 550.

Mertins, Philipp, D. R. Mani, Kelly V. Ruggles, Michael A. Gillette, Karl R. Clauser, Pei Wang, Xianlong Wang, et al. 2016. “Proteogenomics Connects Somatic Mutations to Signalling in Breast Cancer.” Nature; 534 (7605): 55–62.

Michels, Annemieke A., Aaron M. Robitaille, Diane Buczynski-Ruchonnet, Wassim Hodroj, Jaime H. Reina, Michael N. Hall, and Nouria Hernandez. 2010. “mTORC1 Directly Phosphorylates and Regulates Human MAF1.” Molecular and Cellular Biology 30 (15): 3749–57.

Mosca, Roberto, Arnaud Céol, and Patrick Aloy. 2013. “Interactome3D: Adding Structural Details to Protein Networks.” Nature Methods 10 (1): 47–53.

Neale, Benjamin M., Yan Kou, Li Liu, Avi Ma’ayan, Kaitlin E. Samocha, Aniko Sabo, Chiao-Feng Lin, et al. 2012. “Patterns and Rates of Exonic de Novo Mutations in Autism Spectrum Disorders.” Nature 485 (7397): 242–45.

Needham, Elise J., Benjamin L. Parker, Timur Burykin, David E. James, and Sean J. Humphrey. 2019. “Illuminating the Dark Phosphoproteome.” Science Signaling 12 (565). https://doi.org/10.1126/scisignal.aau8645.

Nishi, Hafumi, Kosuke Hashimoto, and Anna R. Panchenko. 2011. “Phosphorylation in Protein-Protein Binding: Effect on Stability and Function.” Structure 19 (12): 1807–15.

Ochoa, David, Mindaugas Jonikas, Robert T. Lawrence, Bachir El Debs, Joel Selkrig, Athanasios Typas, Judit Villen, Silvia Santos, and Pedro Beltrao. 2016. “An Atlas of Human Kinase Regulation. https://doi.org/10.1101/067900.

Olsen, Jesper V., Blagoy Blagoev, Florian Gnad, Boris Macek, Chanchal Kumar, Peter Mortensen, and Matthias Mann. 2006. “Global, in Vivo, and Site-Specific Phosphorylation Dynamics in Signaling Networks.” Cell 127 (3): 635–48.

Paradis, Emmanuel, and Klaus Schliep. 2018. “Ape 5.0: An Environment for Modern Phylogenetics and Evolutionary Analyses in R.” Bioinformatics, July. https://doi.org/10.1093/bioinformatics/bty633.

Pieper, Ursula, Benjamin M. Webb, Guang Qiang Dong, Dina Schneidman-Duhovny, Hao Fan, Seung Joong Kim, Natalia Khuri, et al. 2014. “ModBase, a Database of Annotated Comparative Protein Structure Models and Associated Resources.” Nucleic Acids Research 42 (Database issue): D336–46.

Pollastri, Gianluca, Pierre Baldi, Pietro Fariselli, and Rita Casadio. 2002. “Prediction of Coordination Number and Relative Solvent Accessibility in Proteins.” Proteins 47 (2): 142–53.

Pollastri, Gianluca, Darisz Przybylski, Burkhard Rost, and Pierre Baldi. 2002. “Improving the Prediction of Protein Secondary Structure in Three and Eight Classes Using Recurrent Neural Networks and Profiles.” Proteins 47 (2): 228–35.

Rajesh, Sundaresan, Ružica Bago, Elena Odintsova, Gayrat Muratov, Gouri Baldwin, Pooja Sridhar, Sandya Rajesh, Michael Overduin, and Fedor Berditchevski. 2011. “Binding to Syntenin-1 Protein Defines a New Mode of Ubiquitin-Based Interactions Regulated by Phosphorylation.” The Journal of Biological Chemistry 286 (45): 39606–14.

Ran, F. Ann, Patrick D. Hsu, Jason Wright, Vineeta Agarwala, David A. Scott, and Feng Zhang. 2013. “Genome Engineering Using the CRISPR-Cas9 System.” Nature Protocols 8 (11): 2281–2308.

Reimand, Jüri, Omar Wagih, and Gary D. Bader. 2015. “Evolutionary Constraint and Disease Associations of Post-Translational Modification Sites in Human Genomes.” PLoS Genetics 11 (1): e1004919.

Seringhaus, Michael, Alberto Paccanaro, Anthony Borneman, Michael Snyder, and Mark Gerstein. 2006. “Predicting Essential Genes in Fungal Genomes.” Genome Research 16 (9): 1126–35.

Sharma, Kirti, Rochelle C. J. D’Souza, Stefka Tyanova, Christoph Schaab, Jacek R. Wiśniewski, Jürgen Cox, and Matthias Mann. 2014. “Ultradeep Human Phosphoproteome Reveals a Distinct Regulatory Nature of Tyr and Ser/Thr-Based Signaling.” Cell Reports 8 (5): 1583–94.

Shibano, Takashi, Hiroshi Mamada, Fumihiko Hakuno, Shin-Ichiro Takahashi, and Masanori Taira. 2015. “The Inner Nuclear Membrane Protein Nemp1 Is a New Type of RanGTP-Binding Protein in Eukaryotes.” PloS One 10 (5): e0127271.

Šoštaric, Nikolina, Francis J. O’Reilly, Piero Giansanti, Albert J. R. Heck, Anne-Claude Gavin, and Vera van Noort. 2018. “Effects of Acetylation and Phosphorylation on Subunit Interactions in Three Large Eukaryotic Complexes.” Molecular & Cellular Proteomics: MCP 17 (12): 2387–2401.

Strumillo, Marta J., Michaela Oplova, Cristina Vieitez, David Ochoa, Mohammed Shahraz, Bede P. Busby, Richelle Sopko, et al. 2018. “Conserved Phosphorylation Hotspots in Eukaryotic Protein Domain Families.” https://doi.org/10.1101/391185.

Studer, Romain A., Pascal-Antoine Christin, Mark A. Williams, and Christine A. Orengo. 2014. “Stability-Activity Tradeoffs Constrain the Adaptive Evolution of RubisCO.” Proceedings of the National Academy of Sciences of the United States of America 111 (6): 2223–28.

Studer, Romain A., Ricard A. Rodriguez-Mias, Kelsey M. Haas, Joanne I. Hsu, Cristina Viéitez, Carme Solé, Danielle L. Swaney, et al. 2016. “Evolution of Protein Phosphorylation across 18 Fungal Species.” Science 354 (6309): 229–32.

Teo, Guoci, Guomin Liu, Jianping Zhang, Alexey I. Nesvizhskii, Anne-Claude Gingras, and Hyungwon Choi. 2014. “SAINTexpress: Improvements and Additional Features in Significance Analysis of INTeractome Software.” Journal of Proteomics 100 (April): 37–43.

Torkamani, Ali, Natarajan Kannan, Susan S. Taylor, and Nicholas J. Schork. 2008. “Congenital Disease SNPs Target Lineage Specific Structural Elements in Protein Kinases.” Proceedings of the National Academy of Sciences of the United States of America 105 (26): 9011–16.

Tuoc, Tran Cong, Susann Boretius, Stephen N. Sansom, Mara-Elena Pitulescu, Jens Frahm, Frederick J. Livesey, and Anastassia Stoykova. 2013. “Chromatin Regulation by BAF170 Controls Cerebral Cortical Size and Thickness.” Developmental Cell 25 (3): 256–69.

Türei, Dénes, Tamás Korcsmáros, and Julio Saez-Rodriguez. 2016. “OmniPath: Guidelines and Gateway for Literature-Curated Signaling Pathway Resources.” Nature Methods 13 (12): 966–67.

Tymoszuk, Piotr, Pornpimol Charoentong, Hubert Hackl, Rita Spilka, Elisabeth Müller-Holzner, Zlatko Trajanoski, Peter Obrist, et al. 2014. “High STAT1 mRNA Levels but Not Its Tyrosine Phosphorylation Are Associated with Macrophage Infiltration and Bad Prognosis in Breast Cancer.” BMC Cancer 14 (April): 257.

Uezu, Akiyoshi, Hirokazu Okada, Hideji Murakoshi, Cosmo D. del Vescovo, Ryohei Yasuda, Dario Diviani, and Scott H. Soderling. 2012. “Modified SH2 Domain to Phototrap and Identify Phosphotyrosine Proteins from Subcellular Sites within Cells.” Proceedings of the National Academy of Sciences of the United States of America 109 (43): E2929–38.

UniProt Consortium, The. 2018. “UniProt: The Universal Protein Knowledgebase.” Nucleic Acids Research 46 (5): 2699.

Vaser, Robert, Swarnaseetha Adusumalli, Sim Ngak Leng, Mile Sikic, and Pauline C. Ng. 2016. “SIFT Missense Predictions for Genomes.” Nature Protocols 11 (1): 1–9.

Vizcaíno, Juan Antonio, Attila Csordas, Noemi Del-Toro, José A. Dianes, Johannes Griss, Ilias Lavidas, Gerhard Mayer, et al. 2016. “2016 Update of the PRIDE Database and Its Related Tools.” Nucleic Acids Research 44 (22): 11033.

Wagih, Omar, Marco Galardini, Bede P. Busby, Danish Memon, Athanasios Typas, and Pedro Beltrao. 2018. “A Resource of Variant Effect Predictions of Single Nucleotide Variants in Model Organisms.” Molecular Systems Biology 14 (12): e8430.

Wang, Mingcong, Christina J. Herrmann, Milan Simonovic, Damian Szklarczyk, and Christian von Mering. 2015. “Version 4.0 of PaxDb: Protein Abundance Data, Integrated across Model Organisms, Tissues, and Cell-Lines.” Proteomics 15 (18): 3163–68.

Wen, Z., Z. Zhong, and J. E. Darnell Jr. 1995. “Maximal Activation of Transcription by Stat1 and Stat3 Requires Both Tyrosine and Serine Phosphorylation.” Cell 82 (2): 241–50.

Westmoreland, Tammy J., Sajith M. Wickramasekara, Andrew Y. Guo, Alice L. Selim, Tiffany S. Winsor, Arno L. Greenleaf, Kimberly L. Blackwell, John A. Olson Jr, Jeffrey R. Marks, and Craig B. Bennett. 2009. “Comparative Genome-Wide Screening Identifies a Conserved Doxorubicin Repair Network That Is Diploid Specific in Saccharomyces Cerevisiae.” PloS One 4 (6): e5830.

Widschwendter, Andreas, Sibylle Tonko-Geymayer, Thomas Welte, Günter Daxenbichler, Christian Marth, and Wolfgang Doppler. 2002. “Prognostic Significance of Signal Transducer and Activator of Transcription 1 Activation in Breast Cancer.” Clinical Cancer Research: An Official Journal of the American Association for Cancer Research 8 (10): 3065–74.

Winzeler, E. A., D. D. Shoemaker, A. Astromoff, H. Liang, K. Anderson, B. Andre, R. Bangham, et al. 1999. “Functional Characterization of the S. Cerevisiae Genome by Gene Deletion and Parallel Analysis.” Science 285 (5429): 901–6.

Worby, Carolyn A., Seema Mattoo, Robert P. Kruger, Lynette B. Corbeil, Antonius Koller, Juan C. Mendez, Bereket Zekarias, Cheri Lazar, and Jack E. Dixon. 2009. “The Fic Domain: Regulation of Cell Signaling by Adenylylation.” Molecular Cell 34 (1): 93–103.

wwPDB consortium. 2019. “Protein Data Bank: The Single Global Archive for 3D Macromolecular Structure Data.” Nucleic Acids Research 47 (D1): D520–28.

Zerbino, Daniel R., Premanand Achuthan, Wasiu Akanni, M. Ridwan Amode, Daniel Barrell, Jyothish Bhai, Konstantinos Billis, et al. 2018. “Ensembl 2018.” Nucleic Acids Research 46 (D1): D754–61.

Zhao, Gang, Xiaoke Zhou, Liqun Wang, Guangtao Li, Hermann Schindelin, and William J. Lennarz. 2007. “Studies on peptide:N-Glycanase-p97 Interaction Suggest That p97 Phosphorylation Modulates Endoplasmic Reticulum-Associated Degradation.” Proceedings of the National Academy of Sciences of the United States of America 104 (21): 8785–90.

